# Leucine Aminopeptidase LyLAP enables lysosomal degradation of membrane proteins

**DOI:** 10.1101/2024.12.13.628212

**Authors:** Aakriti Jain, Isaac Heremans, Gilles Rademaker, Tyler C. Detomasi, Grace A. Hernandez, Justin Zhang, Suprit Gupta, Teresa von Linde, Mike Lange, Martina Spacci, Peter Rohweder, Dashiell Anderson, Y. Rose Citron, James A. Olzmann, David W. Dawson, Charles S. Craik, Guido Bommer, Rushika M. Perera, Roberto Zoncu

## Abstract

Proteolysis of hydrophobic helices is required for complete breakdown of every transmembrane protein trafficked to the lysosome and sustains high rates of endocytosis. However, the lysosomal mechanisms for degrading hydrophobic domains remain unknown. Combining lysosomal proteomics with functional genomic data mining, we identify Lysosomal Leucine Aminopeptidase (LyLAP; formerly Phospholipase B Domain-Containing 1) as the hydrolase most tightly associated with elevated endocytic activity. Untargeted metabolomics and biochemical reconstitution demonstrate that LyLAP is not a phospholipase, but a processive monoaminopeptidase with strong preference for N-terminal leucine – an activity necessary and sufficient for breakdown of hydrophobic transmembrane domains. LyLAP is upregulated in pancreatic ductal adenocarcinoma (PDA), which relies on macropinocytosis for nutrient uptake, and its ablation led to buildup of undigested hydrophobic peptides, which compromised lysosomal membrane integrity and inhibited PDA cell growth. Thus, LyLAP enables lysosomal degradation of membrane proteins, and may represent a vulnerability in highly endocytic cancer cells.

**One sentence summary:** LyLAP degrades transmembrane proteins to sustain high endocytosis and lysosomal membrane stability in pancreatic cancer.

## Main Text

Transmembrane proteins represent ∼20-30% of the human proteome and play many essential roles such as nutrient import, signal transduction, cell adhesion and migration (*1–4*). Most transmembrane proteins are subjected to turnover by endocytic uptake and delivery to the lysosome (*5*, *6*), where they are proteolyzed to single amino acids that are ultimately exported to the cytosol (*7*, *8*). Lysosomal degradation of transmembrane proteins serves multiple purposes, including termination of signaling by activated receptor tyrosine kinases, remodeling of inter-cellular and cell-matrix contacts, downregulation of nutrient import (*5*, *9–13*).

In the lysosomal lumen, a set of endo-and exopeptidases recognize substrate proteins via broad and often overlapping target sequences (*14*). Degradation of membrane proteins poses a unique challenge due to their hydrophobic, phospholipid-embedded α-helical domains that are inaccessible to endopeptidases (*7*). While the substrate specificity of most lysosomal proteases, such as cathepsins, has been determined to considerable precision, no lysosomal enzymes that can degrade the hydrophobic α-helical domains spanning the lipid bilayer has been unambiguously identified.

Membrane protein turnover can be greatly accelerated by ligand-stimulated endocytosis, as well as bulk uptake processes, such as phagocytosis, which mediates pathogen clearance by immune cells, and macropinocytosis, a route for nutrient uptake in many cancer types. A notable example is pancreatic ductal adenocarcinoma (PDA), a highly aggressive malignancy that relies on enhanced lysosomal biogenesis and activity to grow in nutrient-poor microenvironments (*15–17*). A feature of PDA is elevated influx of extra-and intracellular macromolecular substrates that reach the lysosome through macropinocytosis and autophagy, respectively (*15*, *16*, *18*). The high macropinocytic flux of PDA cells result not only in increased uptake of soluble extracellular protein cargo, but also in endocytic import of membrane proteins that are not efficiently recycled back to the plasma membrane (*19*, *20*). Whether specific proteins enable PDA lysosomes to handle the enhanced proteolytic load associated with elevated macropinocytosis is unknown.

Endocytic membrane traffic is increasingly linked to lysosomal membrane permeabilization (*21–27*), a process where lipid packing defects in the lysosomal limiting membrane develop into tears that lead to the leakage of luminal contents into the cytosol, triggering various forms of cell death (*28*, *29*). To counter damage, lysosomes rely on an array of repair mechanisms, which engage the lysosomal membrane from the cytoplasmic side (*28*, *29*). For example, a prominent repair mechanism is mediated by the Endosomal Sorting Complex Required for Trafficking (ESCRT)-III, subunits of which oligomerize around and reseal nascent tears in the lysosomal membrane (*25*, *26*). Whether membrane-stabilizing mechanisms act from within the lysosome to support high rates of endocytosis and prevent damage is unknown.

Lysosomal damage has been primarily associated with exogenous substrates including amyloids, viral and bacterial proteins, and other undigestible agents that, upon being trafficked to the lysosome, can breach the limiting membrane via mechanical or chemical stress (*28*, *29*). Due to their hydrophobic nature, transmembrane α-helices derived from proteolysis of membrane proteins may also intercalate in the lysosomal membrane and compromise its stability, if not efficiently degraded (*30*). However, which lysosomal hydrolases prevent membrane damage by breaking down potentially toxic protein domains remains unknown.

### LyLAP is enriched in highly endocytic cells and is required for PDA growth

To discover factors that enable lysosomes to sustain high endocytic flux, we engineered a pipeline that combined functional genomics data mining with organelle proteomics and expression-survival analysis in phagocytic immune cells and PDA cells, both of which are highly reliant on lysosomal function (Fig. 1A). First, we extracted a list of genes required for phagocytosis or for targeted degradation of membrane proteins from four independent CRISPR-Cas9 screens and ranked them based on their expression levels in immune cells (Fig. S1A-S1B) (*31–35*).

**Figure 1.**
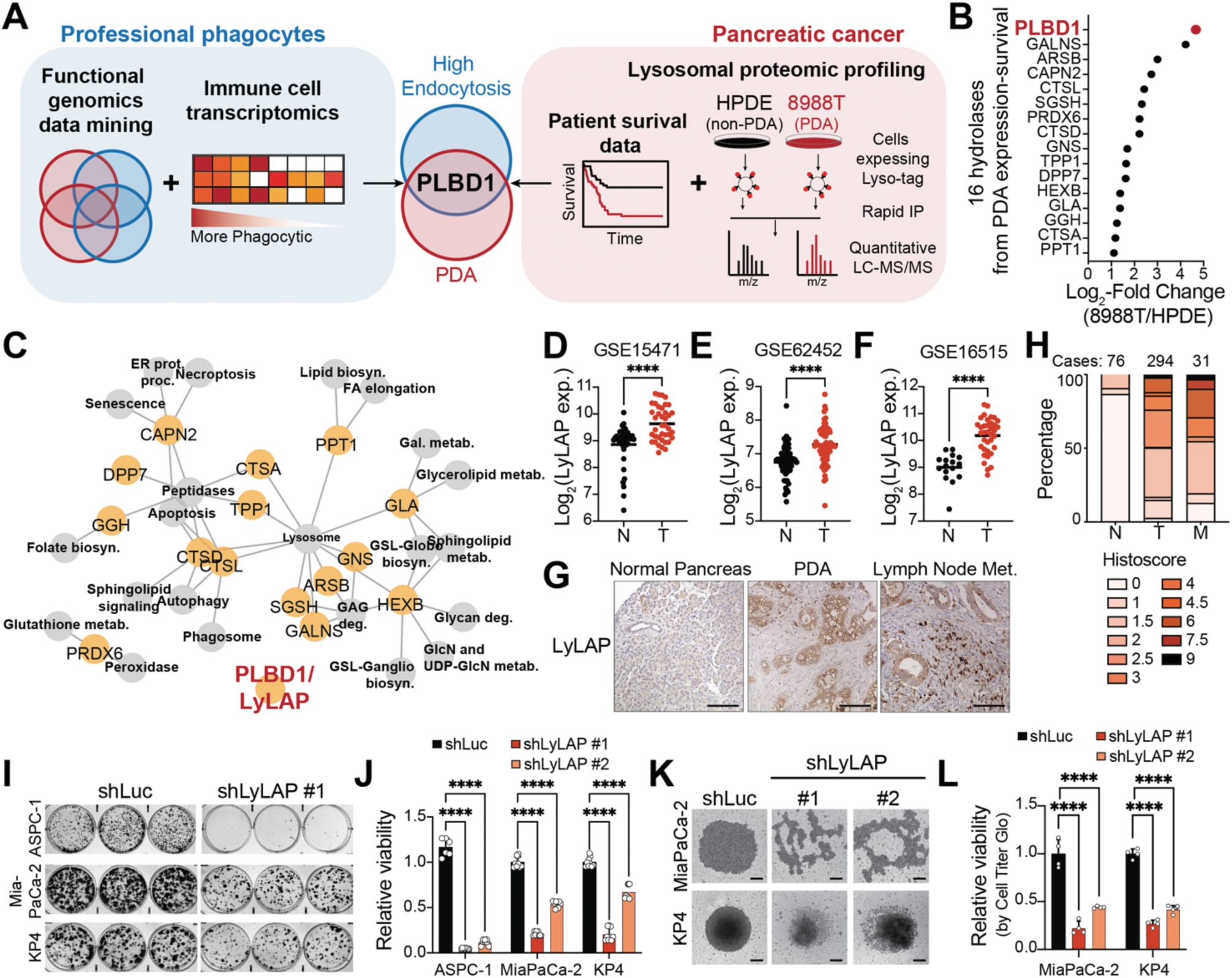
LyLAP is a highly enriched lysosomal hydrolase required for PDA growth. **A.** Summary of bioinformatic pipeline combining functional genomics screens for endocytosis regulators and transcriptomics in immune cells with lysosomal proteomic profiling from 8988T (PDA) versus HPDE (non-PDA) cells and patient survival data from cancer-associated lysosomal hydrolases. B. Log2-fold change in protein abundance from 8988T versus HPDE cells from lysosomal proteomics analysis described in (**A**) of the 16 hydrolases identified by expression and prognostic outcome. C. Network diagram of manually curated functional annotations (from KEGG) of the 16 lysosomal hydrolases identified by expression and prognostic outcome. **D-F.** LyLAP mRNA expression from PDA tumors (T) versus non-tumoral adjacent pancreatic tissue (N) from patient tumor gene expression datasets. Data from Gene Expression Omnibus. **G.** Immunohistochemical staining of LyLAP expression in patient tumor microarrays of normal pancreas (left), PDA (middle), lymph node metastasis (right). Scale bars, 100 μm. **H.** Pathologist-assigned histoscore of LyLAP expression in normal pancreas (N), PDAC tumor (T), and lymph node metastasis (M). **I.** Representative images of colony formation assays from ASPC1, MiaPaCa-2, and KP4 PDA cell lines upon LyLAP knock-down (LyLAP-KD) by shRNA. **J.** Crystal violet dye absorbance from colony formation assay in (**I**) to assess relative viability of ASPC1, MiaPaCa-2 and KP4 cells upon LyLAP-KD. Data normalized to the shLuciferase (shLuc) control of each respective cell line. **K.** Representative images of spheroid formation assay in MiaPaCa-2 and KP4 cells following LyLAP-KD by shRNAs compared to shLuc control. Scale bars, 100 μm. **L.** Quantification of viable cells by CellTiterGlo staining of spheroids in (**K**). Data normalized to the shLuc control of each respective cell line. Data are shown as mean ± SD. Comparison of groups carried out using two-tailed, unpaired t-test (**D-F**) and two-way ANOVA (**J** and **L**). ****p<0.0001

In parallel, we carried out lysosomal immunoisolation and proteomic profiling from Pa-Tu 8988T (PDA) and HPDE (non-transformed counterparts) cells expressing a lysosomal affinity tag (*36–38*), which revealed hydrolases as the most differentially enriched category in PDA versus non-cancer lysosomes (Fig. S1C-S1D). We then ranked PDA-enriched lysosomal hydrolases based on their association with poor prognostic outcome (Fig. 1B and S1E). Remarkably, both the phagocyte-and PDA-focused analyses yielded Phospholipase B Domain-Containing 1 (PLBD1), a putative lysosomal hydrolase, as the top-ranking protein (Fig. 1A, 1B, S1A-S1E). Given the pro-tumorigenic role of the lysosome in PDA, and the prospect of identifying novel druggable targets in this organelle (*39–42*), we decided to further characterize PLBD1 in the context of PDA.

Despite being annotated as a phospholipase, the enzymatic function of PLBD1 has not been firmly established (*43*, *44*). Accordingly, network analysis of the PDA-specific hydrolases based on Kyoto Encyclopedia of Genes and Genomes (KEGG) linked 15 out of the 16 hydrolases to annotated functions but, notably, left PLBD1 unassociated (Fig. 1C). For reasons that will become clear later, we henceforth refer to PLBD1 as Lysosomal Leucine Aminopeptidase (LyLAP).

Across a panel of 1019 cancer cell lines spanning 26 different cancer types, LyLAP had the highest expression score in PDA, with lesser degree of enrichment in other gastrointestinal (GI) tract malignancies (Fig. S1F). In contrast, none of the other 15 hydrolases displayed a comparable enrichment pattern (Fig. S1F). Compared to non-transformed pancreatic cells (HPDE), LyLAP was significantly upregulated in a panel of 11 PDA cell lines by quantitative PCR (qPCR) (Fig. S1G). Consistently, LyLAP was transcriptionally upregulated in patient biopsies compared to normal adjacent tissues (Fig. 1D, 1E, 1F). Moreover, immunohistochemical analysis in patient-derived tumor microarrays showed increased LyLAP expression in primary (ductal) tumor tissue, as well as in lymph node metastases compared to normal pancreatic tissues (Fig. 1G, 1H). As mentioned above, elevated LyLAP expression was associated with poor survival of PDA patients (Fig. S1H).

The elevated expression and negative survival correlation of LyLAP suggests a role for this protein in PDA growth. To test this possibility, we depleted LyLAP using two independent small-hairpin RNAs (shRNAs) that achieved up to 95% knockdown at the mRNA level (Fig. S2A) and undetectable protein levels in lysosomal immunoprecipitates (Fig. S2B). LyLAP depletion in multiple PDA lines led to pronounced growth inhibition in both 2D and 3D growth assays (Fig. 1I-1L, S2C). In contrast, non-transformed pancreatic cells as well as commonly used immortalized cell lines, both of which express low levels of LyLAP, were not significantly affected by LyLAP-targeting shRNAs (Fig. S1H). Thus, elevated expression of LyLAP predicts its requirement for PDA growth.

### LyLAP loss leads to lysosomal dysfunction in PDA

To determine the basis of PDA growth inhibition upon LyLAP depletion, and given its putative role as a lysosomal hydrolase, we examined structural and functional features of lysosomes in LyLAP-depleted cells. Transmission electron microscopy analysis of 3 independent PDA cell lines treated with LyLAP-targeting shRNAs showed the presence of multiple, grossly enlarged (between 4-to 10-fold larger diameter) lysosomal vesicles, with clear disruption of their internal structure (Fig. 2A-2C). In contrast, non-transformed HPDE cells had no discernible changes in lysosomal morphology upon LyLAP knock-down (Fig. 2D). Supporting the ultrastructural analysis, immunofluorescence staining for the lysosomal marker LAMP2 showed significant lysosomal enlargement (10-to 25-fold larger area) in PDA (KP4) cell lines depleted for LyLAP, but not in control (HPDE) cells (Fig. S3A, S3B).

**Figure 2.**
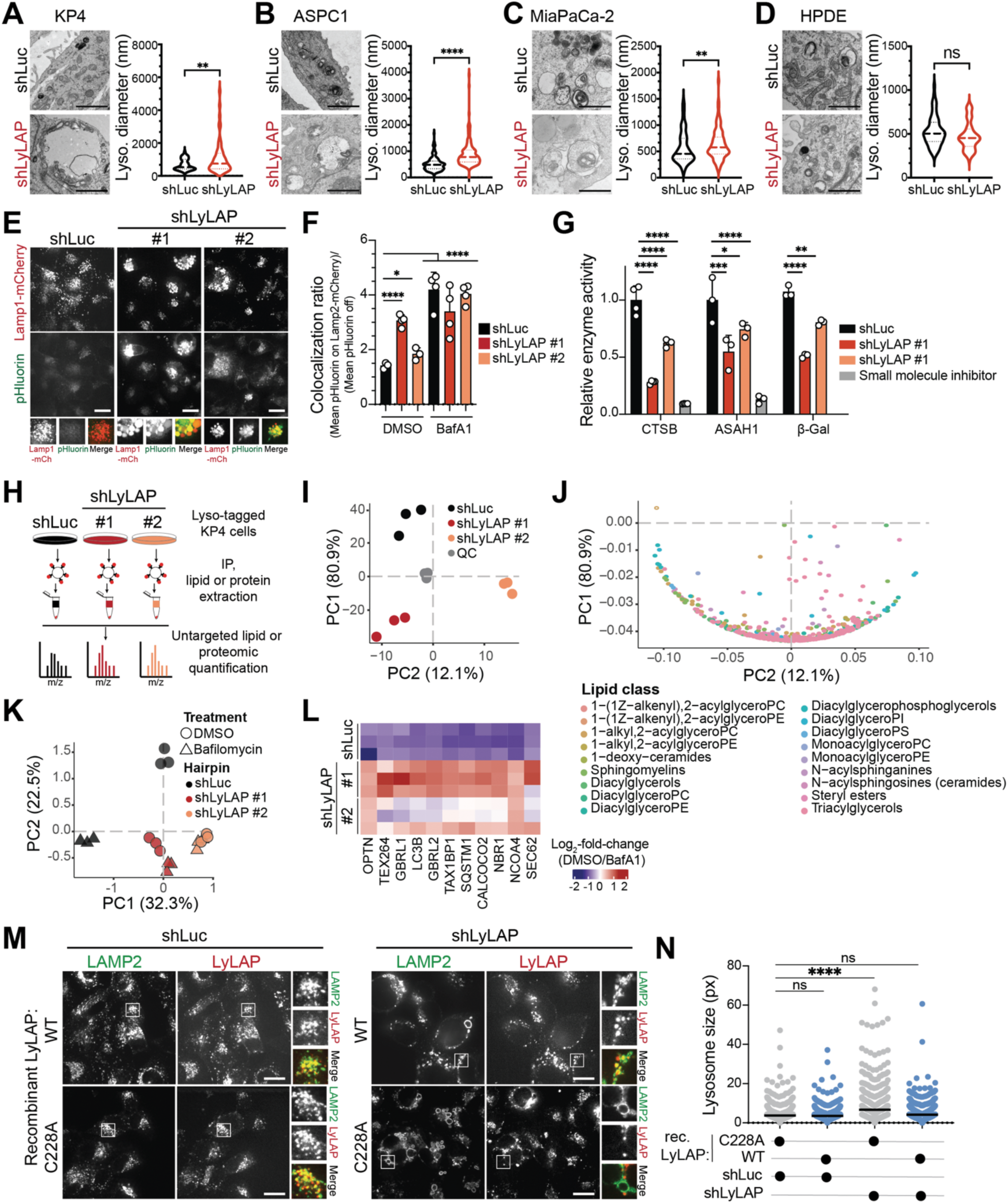
LyLAP depletion causes lysosomal dysfunction in PDA. **A-D.** Representative transmission electron microscopy images of lysosomes from PDA cell lines KP4 (**A**), ASPC1 (**B**), and MiaPaCa-2 (**C**) and the non-transformed control cell line HPDE (**D**) following treatment with shLyLAP or shLuc. Quantification of lysosomal diameter to the right of each respective image. Scale bars, 1 μm. **E.** Representative confocal images of KP4 cells expressing pHluorin-mCherry-LAMP1 following treatment with the indicated shRNAs. Scale bars, 10 μm. **F.** Quantification of relative lysosomal deacidification from (**E**) expressed as colocalization ratio between pHluorin and mCherry. Cells were treated with either DMSO (vehicle control) or Bafilomycin A1 (BafA1, 500 nM) to induce lysosomal deacidification. **G.** Activity assays for Cathepsin B (CTSB), Acid ceramidase (ASAH1), and β-galactosidase (β-Gal) KP4 cells following treatment with the indicated shRNAs. Cells were additionally treated with small molecule inhibitors, Leupeptin/Pepstatin (Leu/Pep) or Carmofur to completely inhibit CTSB and ASAH1, respectively. **H.** Schematic of experimental set-up for untargeted lipidomics and proteomics analysis of lysosomal immunoprecipitates from KP4 cells following treatment with the indicated shRNAs. **I.** Principal component analysis from untargeted lipidomics of KP4 lysosomal immunoprecipitates following treatment with the indicated shRNAs. Quality control (QC) refers to pool of all samples. **J.** Loadings plot of the PCA analysis in (**I**) showing enrichment of multiple lipid species in LyLAP-KD samples. **K.** Principal component analysis from lysosomal proteomics of KP4 lysosomal immunoprecipitates following treatment with the indicated shRNAs. Cells were treated with either DMSO or BafA1 (500 nM) prior to lysosome immunocapture. **L.** Heatmap showing the fold-change in abundance of autophagic adaptors from lysosomes treated with BafA1 versus vehicle (DMSO) prior to immunocapture, from cells treated with the indicated shRNAs. LyLAP-KD resulted in smaller difference between DMSO and BafA1, consistent with impaired baseline proteolysis. **M.** Representative confocal images of KP4 cells treated with indicated shRNAs and supplemented with AF546-maleimide-labeled recombinant LyLAP (WT or C228A) followed by immunofluorescence staining for LAMP2. Scale bars, 10 μm. **N.** CellProfiler-aided quantification of lysosomal size from experiment in (**M**). Data are shown as mean ± SD. Comparison of groups carried out by two-tailed, unpaired t-test (**A-D**), two-way ANOVA (**F, G**), and one-way ANOVA (**N**). ****p<0.0001, ***p<0.001, **p<0.01. *p<0.05, ns=not significant.

A major hallmark of lysosomal dysfunction is loss of luminal acidification established by the vacuolar H^+^-ATPase (*8*, *45*). LyLAP-depleted KP4 cells expressing a genetically encoded lysosomal pH sensor (mCherry-pHluorin-LAMP1; RpH) (*46*) partially lost their lysosomal acidification, as shown by pHluorin unquenching, but to a lesser degree than with the vacuolar H^+^-ATPase inhibitor, Bafilomycin A1 (BafA1) (Fig. 2E, 2F). Most lysosomal hydrolases require acidic pH values (4.5-5) for maximal activity (*8*, *14*). Consistent with loss of lysosomal acidification, the activities of multiple hydrolases, including cathepsin B, acid ceramidase, and β-galactosidase, were partially inhibited in LyLAP-depleted KP4 cells, but to a lesser degree than using specific small-molecule inhibitors of these enzymes (Fig. 2G).

The loss of multiple hydrolase activities suggested that LyLAP-depleted lysosomes should accumulate undegraded substrates. Consistently, lysosome immunopurification followed by untargeted lipidomic analysis revealed buildup of 650 lipid species upon LyLAP depletion (Fig. 2H-2J, S3C, S3D). LyLAP depletion also caused a proteolytic defect, as shown by the accumulation of substrate proteins normally degraded at high rates such as selective autophagy adaptors (*47–49*), although not to the same extent as control lysosomes following treatment with BafA1 (Fig. 2H, 2K, 2L, S3E).

Thus, the general loss of hydrolase activity within LyLAP-depleted lysosomes is accompanied by the buildup of both lipid and protein substrates, providing a rationale for the enlarged lysosomal morphology.

### LyLAP is proteolytically activated within lysosomes

To identify the enzymatic function of LyLAP, we generated recombinant human LyLAP from HEK293 GnTi-cells (Fig. S4A). A crystal structure of bovine PLBD1/LyLAP classified it as an N-terminal nucleophile (Ntn) hydrolase (*43*). All Ntn-hydrolases harbor a cysteine, serine, or threonine nucleophilic residue, which cleaves amide bond-containing substrates. The same residue also mediates autocatalytic maturation of the proenzyme, by cleaving the peptide bond immediately preceding the nucleophile, resulting, in the case of LyLAP, in two domains that remain electrostatically bound (*43*, *50*). Consistent with LyLAP being a lysosomal Ntn-hydrolase, the recombinant 65 kDa purified protein autocatalytically cleaved itself into an N-terminal 26.9 kDa segment and a C-terminal 43 kDa segment (Fig. S4B). Notably, autocatalytic cleavage only occurred when LyLAP was incubated at pH 4.5, strongly suggesting that this maturation step follows proenzyme delivery to the lysosome. Based on homology to the bovine structure, C228 is the nucleophilic residue in human LyLAP and, accordingly, mutating C228 to Ala blocked autocatalytic cleavage irrespective of pH (Fig. S4B).

To test whether self-activated recombinant LyLAP is functional in its native environment, we labeled it with Alexa Fluor 546 (AF546)-maleimide and added it to the media of KP4 cells, which led to LyLAP uptake and delivery to lysosomes (Fig. S4C). Recombinant wild-type LyLAP partially rescued proliferation and colony formation ability of KP4 cells treated with LyLAP-targeting shRNAs. In contrast, the catalytically dead C228A mutant failed to rescue any of the growth defects (Fig. S4D-S4F). Recombinant wild-type LyLAP, but not the C228A mutant, also rescued lysosomal enlargement caused by depletion of endogenous LyLAP (Fig. 2M, 2N). Together, these data strongly indicate that recombinantly expressed LyLAP is active when delivered to lysosomes.

To gain a complete picture of the mechanism of LyLAP activation, we compared the bovine crystal structure with the AlphaFold prediction for full-length human LyLAP (*51*). This comparison highlighted a 19 amino acid segment (L209-H227) in the predicted full-length protein, located proximal to catalytic C228, which was missing from the mature bovine structure (Fig. S4G). We reasoned that removal of this segment via a second cleavage at its N-terminal end (L209) is required to fully activate LyLAP.

Based on precise molecular mass determination by MALDI-TOF, combined with structural considerations we ruled out that C228 cleaved distally at L209 (see Supplementary Text and Fig. S4H-S4J). Moreover, because addition of exogenous, recombinant LyLAP to KP4 cells resulted in rescue of lysosomal morphology and partial rescue of cell growth (Fig. 2M, 2N, S4E, S4F), we reasoned that the L209 position in LyLAP is cleaved by a distinct lysosomal protease. Confirming this hypothesis, when we reconstituted lysosomal immunoprecipitates from KP4 cells with self-cleavage incompetent LyLAP-C228A, we observed the production of the mature N-terminal domain of the expected molecular size (∼24 kDa) (Fig. S4K-S4M). This activation step was inhibited by neutral pH and by an inhibitor of Cys/Ser/Thr cathepsin proteases, leupeptin, but not by aspartyl protease inhibitors, pepstatin or E64D (Fig. S4L, S4M). Based on this result, we tested a small panel of Cys/Ser/Thr cathepsins and found that both Leupeptin-sensitive cathepsin K and L could cleave LyLAP-C228A, whereas cathepsin B and D did not (Fig. S4N, S4O). Further confirming the dual cleavage pattern, incubation with cathepsin L shortened the N-terminal domain created by the autocatalytic cleavage of the wild-type LyLAP (Fig. S4P, S4Q); this fully active, mature form of wild-type LyLAP will henceforth be referred to as LyLAP^active^, whereas the cathepsin K/L-treated, catalytically inactive C228A mutant will be referred to as LyLAP^dead^.

Having obtained fully mature LyLAP^active^, we tested whether it has phospholipase B activity, as suggested by its annotation (*44*). Incubating LyLAP^active^ with C16:0/C18:1 phosphatidylcholine (16:0/18:1 PC) yielded no decrease in 16:0/18:1 PC substrate, or detectable production of the associated lysophosphatidylcholine products, irrespective of pH, as assessed by LC-MS (Fig. S5A, S5B). Because Ntn-hydrolases are amidases, we next tested whether LyLAP^active^ could degrade amide bond-containing lipids found in lysosomes (*50*, *52*). However, LyLAP^active^ had no detectable activity against ceramides (Fig. S5C, S5D), glycosphingolipids, including glucosylceramide (Fig. S5E, S5F), galactosylceramide, and gangliosides (GM3, GD3, AsialoGM1, GM1, GD1a, GD1b, GT1b) (Fig. S5G-S5L), or bile salts (Fig. S5M, S5N).

### LyLAP is a processive monoaminopeptidase for hydrophobic amino acids

The previous data show that LyLAP is unlikely to be a lipid hydrolase. Thus, we next searched for putative non-lipid substrates using untargeted polar metabolomics (*53*). This analysis revealed the statistically significant accumulation of 264 metabolite features in lysosomes upon LyLAP knockdown (Fig. 3A, S6A, S6B). The large number of accumulated metabolites was consistent with the general lysosomal dysfunction upon LyLAP depletion (Fig. 2). To distinguish direct substrates from secondarily accumulated species, we reconstituted LyLAP-depleted lysosomal lysates with LyLAP^active^ or LyLAP^dead^ (as well as cathepsin L alone) (Fig. 3B). We then looked for species that accumulate upon LyLAP loss and are specifically depleted by recombinant LyLAP^active^add-back, but not under any other condition.

**Figure 3.**
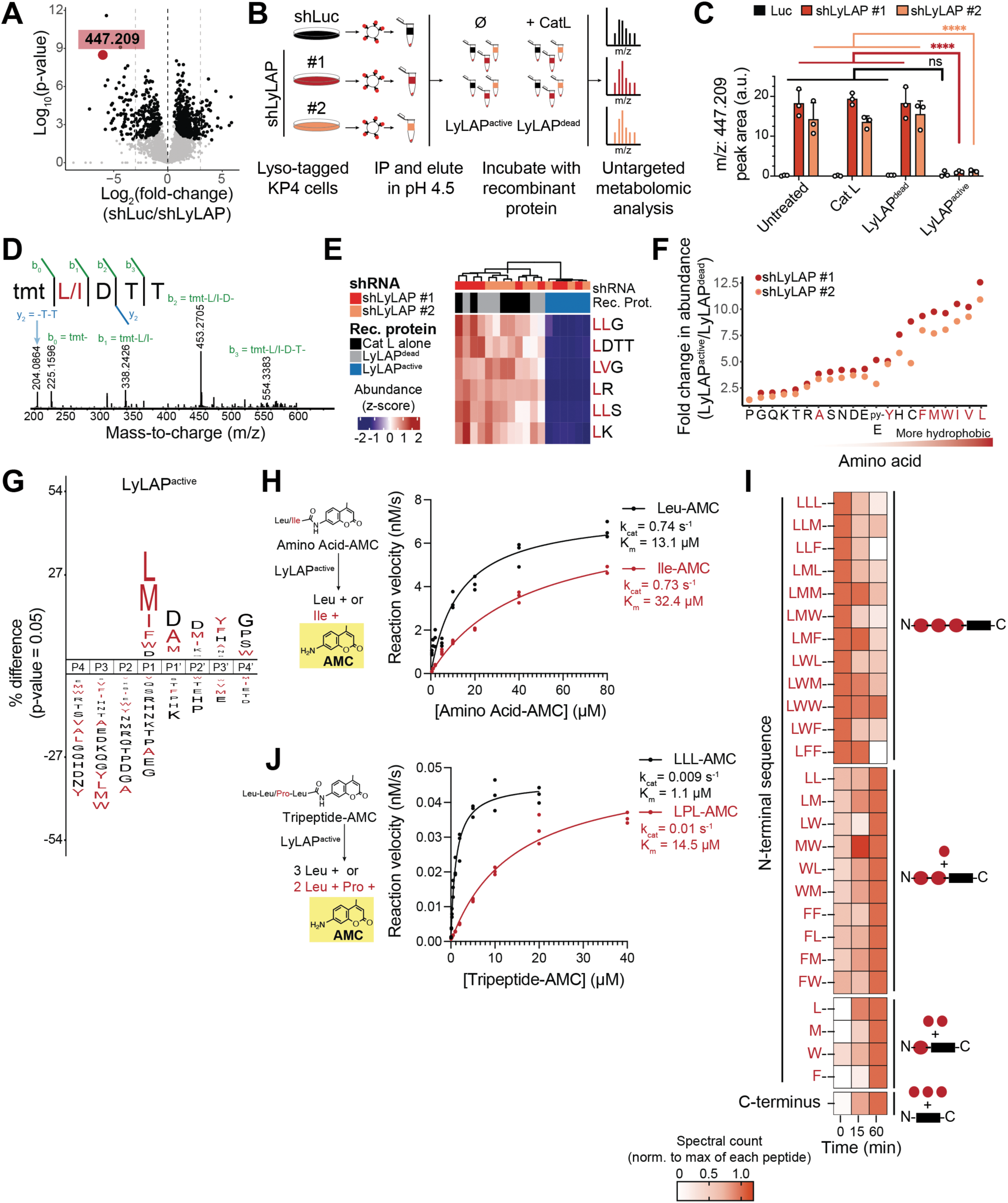
LyLAP is a processive monoaminopeptidase for hydrophobic amino acids. **A.** Volcano plot from untargeted metabolomic analysis of lysosomes immunoprecipitated from KP4 cells treated with shLuc or shLyLAP. Highlighted m/z ratio is a putative substrate. **B.** Schematic showing the experimental set-up for untargeted metabolomic analysis of shLuc or shLyLAP KP4-derived lysosomal immunoprecipitates, following reconstitution with recombinant LyLAP^active^, LyLAP^dead^, or Cathepsin L alone. **C.** Quantification of m/z=447.209 following untargeted metabolomic analysis from (**B**). Data are shown as mean ± SD. Comparison of groups carried out by two-way ANOVA. ****p<0.0001, ns=not significant. **D.** MS2 spectra of the peptide corresponding to TMT-derivatized m/z=447.209. **E.** Heatmap of metabolites detected in experiment described in (**B**) that accumulate upon LyLAP-KD and are only depleted upon incubation with recombinant LyLAP^active^. Metabolites are identified as peptides based on MS2 spectra from **Fig. S6E-S6I**. **F.** Fold-change in abundance of single amino acids shLyLAP KP4-derived lysosomal immunoprecipitates, following reconstitution with recombinant LyLAP^active^ or LyLAP^dead^. Hydrophobic amino acids are annotated in red. **G.** ICE-logo from MSP-MS data for LyLAP^active^. Letters above the midline indicate preference for residue at that position for cleavage (positions P4-P4’ shown), letter height indicates larger preference for that residue. Pattern where positions P4-P2 are not preferred is indicative of a monoaminopeptidase. Logo represents cleavages from technical triplicates and peptide comparison to all possible cleavages in the MSP-MS library and weighted by spectral count observed across all timepoints. **H.** (*left*) Schematic of aminopeptidase activity of LyLAP^active^ determined by incubation with appropriate amino acid (AA) conjugated to AMC as substrate, which is cleaved to free amino acid and fluorescent AMC as products. (*right*) Michaelis-Menten curves of LyLAP^active^ activity against Leu-AMC substrate (black) and Ile-AMC substrate (red). **I.** Heatmap of normalized peptide spectral counts over time from a synthetic peptide library designed to evaluate processive monoaminopeptidase cleavage activity of LyLAP^active^. C-terminus refers to the following peptide sequence: KAHSDVWPYQDA. **J.** (*left*) Schematic of processive aminopeptidase activity of LyLAP^active^ determined by incubation with appropriate tripeptide conjugated to AMC as substrate, which is proteolyzed to free amino acid constituents and fluorescent AMC as products. (*right*) Michaelis-Menten curves of LyLAP^active^ activity against LLL-AMC tripeptide substrate (black) and LPL-AMC tripeptide substrate (red).

Out of the 264 accumulated features, a single species with mass-to-charge (m/z) ratio 447.209, followed this pattern (Fig. 3C). Within the same analysis, we noted along with the disappearance of the 447.209 species, the accumulation of multiple free amino acids in the LyLAP^active^ -incubated samples (Fig. S6C, S6D). This observation favored the assignment of the m/z = 447.209 species as a peptide. Tandem mass tagging (TMT) followed by MS/MS analysis of the same samples identified the m/z = 447.209 species as either Leu-Asp-Thr-Thr (LDTT) or Ile-Asp-Thr-Thr (IDTT) (Fig. 3D). The TMT-MS/MS analysis identified additional peptide species that were specifically degraded by LyLAP^active^ (Fig. 3E, S6E-S6I). Notably, all the LyLAP-degraded peptides had either Leu or Ile at their N-termini (Fig. 3D, 3E, S6E-S6I).

We noted that Leu and Ile, found at the N-termini of the candidate substrate peptides, were also among the most enriched free amino acids upon treatment of the lysosomal lysates with LyLAP^active^ (Fig. 3F). This suggested that LyLAP may be a monoaminopeptidase that selectively cleaves hydrophobic amino acids from intermediate peptide products of lysosomal proteolysis. To independently test this hypothesis, we challenged recombinant LyLAP^active^ with a synthetic peptide library with sufficient complexity to identify consensus cleavage sites for both endo-and exopeptidases (*54*). Multiplexed substrate profiling-mass spectrometry (MSP-MS) of the resulting cleavage products unequivocally identified hydrophobic N-terminal amino acids, especially leucine, as the preferred cleavage sites for LyLAP, strongly supporting its hydrophobic-directed monoaminopeptidase function (Fig. 3G, S7A).

To determine the kinetic parameters of LyLAP, we carried out an *in vitro* cleavage assay, incubating LyLAP^active^ (and appropriate controls) with various amino acids C-terminally conjugated to the fluorogenic substrate, 7-amino 4-methyl coumarin (AMC), which is unquenched upon cleavage (Fig. 3H). LyLAP^active^ exhibited aminopeptidase activity against Leu-AMC and Ile-AMC, with k_cat_ = 0.74 s^-1^ and 0.73 s^-1^, respectively, lower K_m_ for Leu versus Ile (13.1 μM and 32.4 μM, respectively), consistent with the MSP-MS analysis, at a pH optimum of 4 (Fig. 3H, S7B-S7D). Consistent with the untargeted metabolomics result (Fig. 3F), LyLAP did not cleave Pro-AMC and Gly-AMC (S7E, S7F).

Several LyLAP peptide substrates from the untargeted metabolomics (Fig. 3E, S6E-S6G) and MSP-MS analysis (Fig. S7A) contained two or more consecutive hydrophobic residues at their N-termini. Thus, we next asked whether the monoaminopeptidase activity of LyLAP could be processive, enabling complete degradation of hydrophobic N-terminal stretches in substrate peptides. We challenged recombinant LyLAP^active^ with a synthetic 15-mer peptide library, in which the N-terminal Leu is followed by two residues comprising combinations of hydrophobic amino acids (Leu, Met, Phe, and Trp), followed by a non-hydrophobic 12-amino acid peptide sequence. Consistent with processive monoaminopeptidase activity, we observed time-dependent shortening of the 15-mers down to their LyLAP-resistant 12-amino acid C-termini, and a corresponding increase in partially degraded 14-and 13-mer peptides (Fig. 3I). To determine the kinetic parameters of processive LyLAP activity, we carried out an *in vitro* cleavage assay with tripeptides C-terminally conjugated to AMC. LyLAP^active^ exhibited processive aminopeptidase activity against Leu-Leu-Leu-AMC and Leu-Pro-Leu-AMC, with kcat = 0.009 s^-1^ and 0.01 s^-1^ and Km = 1.1 μM and 14.5 μM, respectively, indicating strong preference for higher Leu content (Fig. 3J, S7G, S7H).

Finally, comparison of LyLAP^active^ with the non-Cathepsin L-processed, but self-cleaved enzyme, showed that removal of the L209-H227 loop by Cathepsin K/L increased catalysis toward both single amino acid substrates and, more appreciably, toward peptide substrates (Fig. S7B, S7C, S7G, S7H).

Together, these data strongly support a processive, Leu-directed monoaminopeptidase activity for LyLAP.

### LyLAP enables transmembrane protein degradation in PDA lysosomes

The previous results indicated that LyLAP can remove N-terminal hydrophobic residues from peptides in a processive manner *in vitro*. To identify the endogenous peptide substrates degraded by LyLAP within lysosomes, we supplemented LyLAP-depleted lysosomal lysates with the recombinant, active or inactive, enzyme followed by peptidomic analysis (Fig. 4A, S8A, see Supplemental Text). This analysis confirmed that the endogenous peptide substrates of LyLAP matched the consensus identified by the synthetic MSP-MS library (Fig. 4B, S8B). Moreover, through a multi-parametric analysis of peptide features, the most significant difference was the higher hydrophobicity of LyLAP substrate versus non-substrate (Fig. 4C, S8C). Consistently, the peptides that accumulated in LyLAP-depleted versus control lysosomes also showed higher hydrophobic content (Fig. 4D-4F).

**Figure 4.**
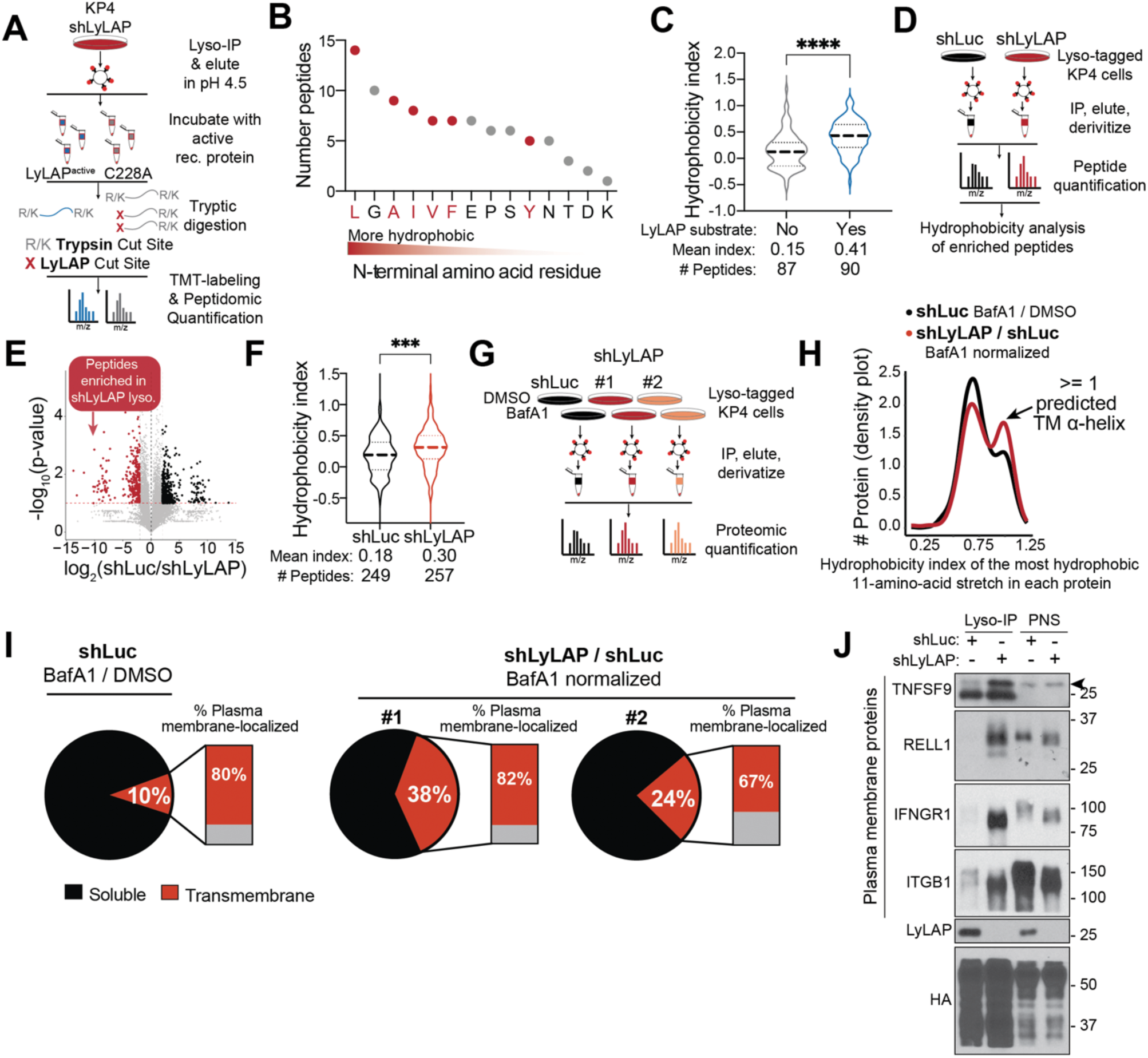
LyLAP enables transmembrane protein degradation in PDA lysosomes. **A.** Schematic of experimental set-up for semi-tryptic peptidomic analysis of LyLAP activity. shLuc or shLyLAP KP4-derived lysosomal immunoprecipitates, were reconstituted with recombinant LyLAP^active^ or LyLAP^dead^, followed by protease digestion with Trypsin and LysC, and quantified by LC-MS. **B.** Frequency of N-terminal amino acid residue on N-semitryptic peptides identified as LyLAP substrates. **C.** Hydrophobicity index (by Abraham-Leo scale) of peptides identified as LyLAP substrates (enriched in LyLAP^dead^ over LyLAP^active^) compared to non-LyLAP substrate peptides. **D.** Schematic of experimental set-up for peptidomic analysis of shLuc or shLyLAP KP4-derived lysosomal immunoprecipitates. **E.** Volcano plot of peptides enriched in lysosomes from shLyLAP KP4 cells versus shLuc cells from experiment described in (**D**). Points annotated in black and red refer to peptides with a p-value < 0.05 and log2-fold-change greater than 2 or less than -2, respectively. **F.** Hydrophobicity index (by Abraham-Leo scale) of peptides enriched in LyLAP-depleted lysosomes compared to control lysosomes. **G.** Schematic of experimental set-up for proteomic analysis of shLuc or shLyLAP KP4-derived lysosomal immunoprecipitates. Samples were treated with either BafA1 or vehicle (DMSO) control prior to lysosome immunoprecipitation, TMT-derivatization, and proteomic quantification. **H.** Density plot of hydrophobicity index, revealing a bimodal distribution where the peak to the right represents proteins with 1 more transmembrane α-helices. Control and LyLAP-depleted samples are indicated in black and red, respectively. **I.** Pie chart indicating the relative proportion of transmembrane proteins to total protein substrates in lysosomal immunoprecipitates following the indicated treatments. Stacked bar graphs represent percent of transmembrane proteins in each group that are plasma membrane localized. **J.** Immunoblot showing defective degradation of multiple plasma membrane derived proteins in KP4 lysosomes following LyLAP depletion. Comparison of groups carried out by two-tailed, unpaired t-test (**C, F**). ****p<0.0001, ***p<0.001.

To relate the hydrophobic peptides cleaved by LyLAP to specific protein substrates, we used lysosomal proteomics to identify undigested proteins that preferentially accumulated in LyLAP-depleted over control lysosomes, when lysosomal proteolysis was shut down using BafA1 (Fig. 4G). The use of BafA1 normalization enabled us to identify and remove LyLAP-independent protein substrates with higher confidence. This analysis showed that, when plotted for hydrophobicity, lysosomal protein substrates are arranged in a bimodal distribution where the second peak represents integral membrane proteins containing one or more transmembrane α-helices (Fig. 4H) (*55*, *56*). Remarkably, the distribution of *bona fide* LyLAP substrates was skewed towards a higher fraction of transmembrane helix-containing proteins (Fig. 4H).

Consistent with the hydrophobicity analysis, proteins annotated as transmembrane made up a much larger (up to 4-fold) fraction of all substrates in LyLAP-depleted over control lysosomes (Fig. 4I). The majority (up to 82%) of transmembrane proteins that were LyLAP substrates are plasma membrane localized (Fig. 4I). Many plasma membrane proteins are endocytosed and trafficked to the lysosome for degradation, including integrins and signaling receptors (*9*, *12*, *57*). Several members of these two classes were among the LyLAP substrates, including TNFSF9, RELL1, IFNGR1, and ITGB1 (Fig. S8D). Through immunoblotting of immunoprecipitated lysosomal samples, we confirmed the accumulation of these proteins in LyLAP-depleted versus control lysosomes (Fig. 4J).

In conclusion, by linking together amino acid-, to peptide-, to protein-level analysis, we discover that the aminopeptidase activity of LyLAP serves to break down hydrophobic helical domains belonging to transmembrane proteins, most of which are plasma membrane-derived.

### Loss of LyLAP triggers lysosomal membrane permeabilization

We next asked whether failure to degrade hydrophobic peptide domains underlies the profound dysfunction that we observed in LyLAP-depleted lysosomes (Fig. 2). Hydrophobic peptides have the propensity to partition into lipid bilayers (*30*, *58–60*). In a biological membrane, such as the lysosomal limiting membrane, this partitioning could disrupt lipid packing and favor permeabilization, a process that can kickstart a downstream cascade of ion leakage, loss of hydrolase activity and, ultimately, cell death (*61*).

Consistent with underlying lysosomal membrane damage, LyLAP-depleted PDA cells displayed lysosomal recruitment of ESCRT-III components, CHMP4B and CHMP1A, which mark sites of lysosomal membrane damage and contribute to its repair (*25*, *26*) (Fig. 5A-5D). In contrast, these markers remained diffuse in the cytoplasm of control cells (Fig. 5A-5D). Furthermore, LyLAP-depleted cells were more sensitive to L-leucyl-L-leucine methyl-ester (LLOMe), an agent that trigger lysosomal membrane permeabilization through its conversion to Leu polymers that are thought to intercalate in the lysosomal membrane (*62*) (Fig. 5A-5D).

**Figure 5.**
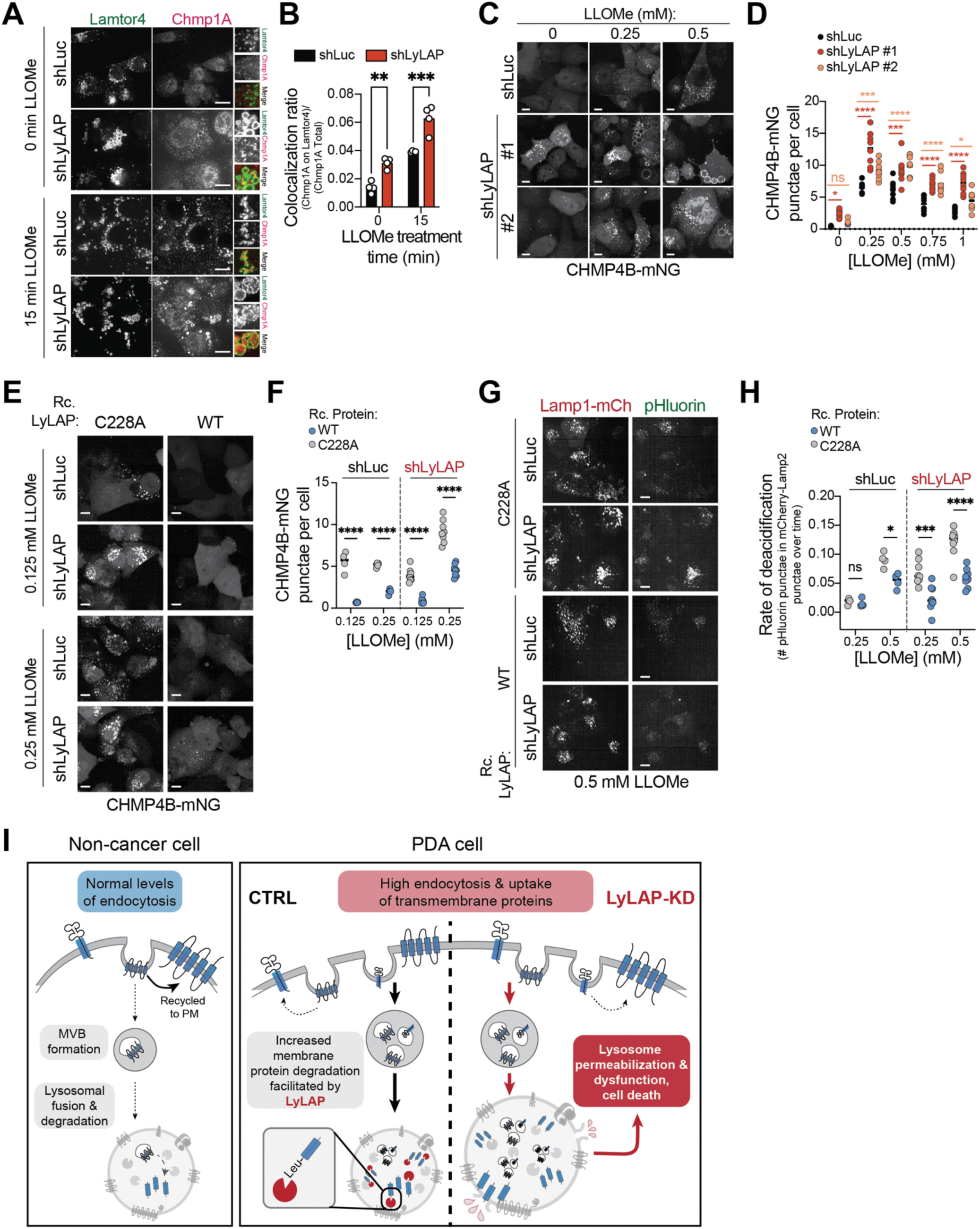
Loss of LyLAP activity triggers lysosomal membrane permeabilization. **A.** Representative immunofluorescence images of KP4 cells transduced with shLyLAP or shLuc and left untreated (*top*) or treated with 0.5 mM LLOMe for 15 minutes (*bottom*). Cells were fixed and stained for Lamtor4 (lysosomal marker) and Chmp1A (ESCRT-III marker). Scale bars, 10 μm. **B.** Quantification of colocalization ratio of Chmp1A and Lamtor4 from immunofluorescence images of shLyLAP-or shLuc-transduced KP4 cells treated with 0.5 mM LLOMe for 15 min. **C.** Representative confocal images of KP4 cells expressing CHMP4B-mNeonGreen (mNG) and transduced with shRNAs against LyLAP (or shLuc control). Cells were treated for 10 minutes with indicated LLOMe concentrations. Scale bars, 10 μm. **D.** Quantification of CHMP4B-mNG punctae per cell upon treatment with the indicated concentrations of LLOMe for 10 minutes. **E.** Representative confocal images of KP4 cells expressing CHMP4B-mNeonGreen (mNG) and transduced with shRNAs against LyLAP (or shLuc control) and supplemented with recombinant C228A or WT LyLAP. Cells were treated for 10 minutes with indicated concentrations of LLOMe. Scale bars, 10 μm **F.** Quantification of CHMP4B-mNG punctae intensity per cell upon treatment with the indicated concentrations of LLOMe for 10 minutes. **G.** Representative confocal images of KP4 cells expressing RpH and transduced with shRNAs against LyLAP (or shLuc control) and supplemented with recombinant C228A or WT LyLAP. Cells were treated with 0.5 mM LLOMe. Lamp1-mCh shown to visualize lysosomes. Scale bars, 10 μm. **H.** Quantification of pHluorin colocalization with Lamp1-mCherry over time upon treatment with the indicated concentrations of LLOMe. **I.** Model illustrating the necessity for LyLAP in highly endocytic PDA cells. Loss of LyLAP activity triggers accumulation of undigested hydrophobic peptides, leading to lysosomal permeabilization and dysfunction, and ultimately cell death. Comparison of groups carried out by two-way ANOVA (**B, D, F, H**). ****p<0.0001, ***p<0.001, **p<0.01, *p<0.05, ns=not significant.

Recombinant wild-type LyLAP, delivered to LyLAP-depleted KP4 cells, was sufficient to rescue the hypersensitivity to LLOMe-induced membrane damage (Fig. 5E, 5F) and deacidification (Fig. 5G, 5H), whereas the inactive C228A showed no protection. Notably, wild-type LyLAP conferred strong protection from LLOMe-induced membrane permeabilization even in control, LyLAP-expressing cells (Fig. 5E, 5F). Combined with the peptide processivity analysis (Fig. 3), this result suggests that excess LyLAP stabilizes LLOMe-challenged lysosomes by directly degrading the LLOMe-derived Leu polymers.

Collectively, these data demonstrate that the monoaminopeptidase activity of LyLAP protects the lysosomal limiting membrane from disruption by undigested hydrophobic peptides. Increased LyLAP levels enable lysosomes to maintain their integrity under high proteolytic loads and may play a stabilizing role in the presence of a variety of chemical and mechanical stressors.

## Discussion

Following their export from the ER, most transmembrane proteins depend on the lysosome for their degradation. Here, we discovered that lysosomes rely on LyLAP to proteolyze the hydrophobic helical domains of membrane protein substrates. Our data indicates that LyLAP is a processive monoaminopeptidase with strong preference for leucine and other hydrophobic residues, often overrepresented in transmembrane domains. LyLAP was highly abundant in lysosomes from PDA cells, enabling degradation of membrane proteins that are likely trafficked in by high endocytic and macropinocytic flux (*63*). Accordingly, inactivating LyLAP resulted in build-up of undigested hydrophobic peptides that disrupted the integrity of PDA lysosomes (Fig. 5I). The discovery of LyLAP fills a major gap in our understanding of membrane protein degradation and turnover within the lysosome.

Several leucine aminopeptidases, known as LAPs, have been described in the cytosol of both eukaryotic and bacterial cells. These enzymes, which play regulatory roles such as peptide processing for antigen presentation and bioactive peptide activation, are metallopeptidases that require divalent cations for their activity (*64*). LyLAP is the first identified lysosomal leucine aminopeptidase. Its Ntn-hydrolase fold distinguishes it from the cytosolic LAPs, but also from other lysosomal proteases, such as cathepsins (*14*). Our data supports the previous hypothesis, based on purely structural grounds, that C228 is the nucleophile that mediates both autocatalytic and substrate cleavage (*43*). Further structural studies of the enzyme-substrate complex will be required to elucidate the details of the catalytic mechanism.

Transmembrane proteins are degraded within the lysosomal lumen following their capture into intraluminal vesicles (ILV), which invaginate from the endosomal limiting membrane (*5*, *6*). Once in the lysosomal lumen, ILVs containing membrane proteins are broken down by deglycosidases, lipases and proteases, although whether these events are sequential or simultaneous is unclear. Because LyLAP attacks the N-termini of transmembrane domains, these must be pre-cleaved and exposed by lysosomal endopeptidases. Moreover, it is likely that the substrate peptide needs to be freed of surrounding lipids before it can be processed by LyLAP. Our data suggests that LyLAP-mediated degradation should occur rapidly to prevent accumulation of undigested hydrophobic domains and the resulting membrane damage. Future studies will be required to reconstitute the chain of events in this pathway.

LyLAP inactivation caused loss of lysosomal membrane integrity, deacidification, partial inactivation of several hydrolases with buildup of their respective substrates, and, ultimately, PDA cell death. A likely mechanism for this toxicity is the partitioning of undigested hydrophobic peptides into the lysosomal limiting membrane, an event that could disrupt lipid packing and cause its permeabilization (*30*, *58–60*). In support of this model, LyLAP-depleted PDA lysosomes had baseline ESCRT-III recruitment and increased sensitivity to LLOMe (Fig. 5). However, the undigested hydrophobic peptides could exert other toxic actions beside membrane intercalation (*29*, *65*).

We discovered LyLAP as the most differentially enriched hydrolase in PDA versus non-cancer lysosomes. Moreover, LyLAP is a highly expressed hydrolase in PDA patients both by gene expression and histochemical analysis. PDA requires high rates of endocytic membrane traffic and autophagy for its growth (*63*). While this traffic benefits PDA growth by providing nutrients from the extracellular environment, it may also deliver large amounts of plasma membrane proteins to the lysosome. Our data are most consistent with the hypothesis that elevated LyLAP expression helps PDA lysosomes to cope with the enhanced proteolytic load caused by endocytic influx. PDA cells were shown to upregulate transcriptional programs for lysosomal biogenesis as an adaptive mechanism to enhance macromolecular breakdown, nutrient scavenging, and stress adaptation (*16*, *18*, *38*, *63*, *66*). Whether LyLAP is part of this broad program or is separately regulated under specific settings remains to be determined.

In support of our model that LyLAP upregulation enables cells to cope with increased endocytic flux, this enzyme is also highly upregulated in professional phagocytic cells, such as macrophages and trophoblasts (Fig. S1B, S9A). We speculate that in the context of pathogen engulfment, a similar influx of transmembrane proteins belonging to either the host or the pathogen might impose a proteolytic burden on macrophage lysosomes, thereby requiring LyLAP activity. A related question is how cells that do not express high levels of LyLAP (which include several non-PDA cancers as well as non-phagocytic tissues) degrade transmembrane proteins in the lysosome. One possibility is that other enzymes with hydrophobic aminopeptidase activity might exist. For example, LyLAP/PLBD1 has a widely expressed paralog, PLBD2 (*67*), which could provide baseline capability for transmembrane protein degradation. Cathepsin H was also shown to N-terminally cleave hydrophobic amino acids *in vitro*, although its processivity and substrate specificity were not determined (*68*). Alternatively, when influx rates of transmembrane proteins are relatively low, lysosomes may dispose of undigested hydrophobic stretches through alternative mechanisms such as intramembrane proteases or lysosomal exocytosis (*8*, *69*).

## Supporting information

Supplemental data combined

## Acknowledgements

The authors thank Dr. C. W. Kombala and Dr. K. Brandvold for sharing fluorogenic bile salt reagents and Dr. M. Stagi and Prof. H. Stenmark for sharing plasmids. We thank Dr. K. Aoki and Prof. M. Tiemeyer for quantification of cellular glycolipid levels and Prof. B. Nagar for consultation on Ntn-hydrolases.

## Author contributions

A.J., R.M.P. and R.Z. conceived the study and designed the experiments. A.J., I.H., G.R., T.C.D, G.H., J.Z., S.G., T.vL., M.L., M.S., P.R. performed experiments and analyzed data. M.L., J.O. and R.C. provided experimental advice. D.A. provided reagents, and D.D. provided clinical samples. G.B. and C.C. supervised the untargeted metabolomic and MSP-MS components of the work, respectively. R.M.P. supervised all aspects related to PDA biology.

A.J. and R.Z. wrote the manuscript with input from all co-authors.

## Data materials and availability

All data are available in the manuscript or supplementary materials or available upon request.

## Supplemental materials

Materials and Methods

Supplementary Text

Figures S1 to S9

Data S1 to S9

## Materials and Methods

### Chemicals and antibodies

Primary antibodies were obtained from the following sources: HA (3724), TOM20 (42406), PDI (3501), GM130 (12480), PEX5 (83020), SQSTM1 (39749), TAX1BP1 (5105), NBR1 (9891), GABARAPL1 (13733), LC3B (3868), and Lamtor4 (13140) from Cell Signaling Technologies; Calcoco2 (12229-1-AP), TNFSF9 (66450-1-Ig), RELL1 (29823-1-AP), IFNGR1 (10808-1-AP), and ITGB1 (12594-1-AP) from Proteintech; LAMP2 (sc-18822), His (sc-57598), and Chmp1a (sc-271617) from Santa Cruz Biotechnology; LyLAP/PLBD1 (HPA040303) from Sigma Aldrich; and NPC1 (ab134113) from Abcam. Secondary antibodies were obtained from the following sources: anti-Ms-AF488 (A11001), anti-Rb-AF488 (A11011), anti-Ms-AF568 (A11004), anti-Rb-AF568 (A11008) from Thermo Fisher; anti-Ms-HRP (A4416) from Sigma Aldrich, anti-Rb-HRP (PI-1000) from Vector Laboratories.

Chemicals were obtained from the following sources: Bafilomycin A1 (J61835) and Leupeptin hemisulfate (J61188) from Alfa Aesar; L-Leucyl-L-Leucine methyl ester (hydrochloride) (LLOMe, 16008) and Gly-Phe-β-naphthylamide (GPN, 14634) from Cayman Chemical; Pepstatin A (195368) from MP Biomedicals; E64D (E8640) from Sigma Aldrich.

### Plasmids and cloning

Short-hairpin oligonucleotide (shRNA) directed against LyLAP/PLBD1 (TRCN0000425993 for shLyLAP #1 and TRCN0000416996 for shLyLAP #2) or Luciferase (TRCN0000072243, used as a non-targeting control) were cloned into the pLKO.1 lentiviral vector (the RNAi Consortium, Broad Institute) according to manufacturer instructions.

Synthetic cDNA encoding TMEM192-RFP-3xHA was sub-cloned into the pLVX lentiviral vector (*37*). Inserts encoding mCherry-pHluorin-Lamp1-3xFlag (RpH) (*46*) and CHMP4B-mNG (*25*) were sub-cloned into the pCDH lentiviral vector. Synthetic cDNA encoding for IgK-TEV-6xHis-PLBD1, IgK-TEV-6xHis-PLBD1(C228A), and IgK-TEV-6xHis-ASAH1 were cloned into the pRK5 vector for mammalian expression.

### Mammalian cell culture

KP4, MiaPaCa-2, ASPC-1, Pa-Tu 8988T, and 293T cells and their derivatives were cultured in DMEM media (Gibco, 11965) with 10% fetal bovine serum supplemented with 1% penicillin and streptomycin. HPDE cells and their derivatives were cultured in defined Keratinocyte SFM media (Gibco, 10744019). All adherent cell lines were maintained at 37°C and 5% CO_2_. Suspension 293 GnTI-cells were cultured in FreeStyle 293 expression medium (Gibco, 12338018) supplemented with 2% FBS and 1% penicillin and streptomycin and maintained at 37°C and 8% CO_2_, while shaking at 130 rpm. All cells were routinely tested for mycoplasma contamination using mycoplasma PCR detection kit (abm, G238).

### Human samples

The tissue microarray containing pancreatic cancer specimens has been detailed in a previous publication (*70*). Human pancreatic ductal adenocarcinoma (PDA) samples were procured from patients who had undergone surgical resection of their primary PDA tumors. The samples were gathered at UCLA in accordance with the approved protocol IRB#11-000512. Notably, no selection biases were identified. The study was carried out in strict adherence to institutional ethical guidelines, with minimal risk as per the IRB protocol, hence formal informed consent was not required.

### Lentiviral production and transduction

Lentiviral particles were generated by co-transfecting pLKO, pLVX or pCDH constructs with pMD2.G (Addgene, 12259) and psPAX2 (Addgene, 12260) packaging plasmids at a 5:3.75:1.25 ratio to HEK293T cells using PEI transfection reagent (Polysciences Inc., 23966). Viral supernatants were collected 48 h and 72 h post-transfection and filtered using a 0.45 μm syringe filter. Virus was concentrated using Lenti-X concentrator (Clontech, 631231) according to manufacturer instructions and stored at -80°C prior to transduction. For lentiviral transduction, the virus and 8 μg/mL polybrene (Millipore TR-1003-G) were added to target cells 24 h after seeding. After 24 h, virus was removed, and cells were selected with fresh media containing either puromycin (1.5 mg/mL), hygromycin B (200 mg/mL), or blasticidin (10 mg/mL). Experiments were performed 4-6 days post transduction.

### Spheroid growth assay

2000 KP4 or MiaPaCa-2 cells (transduced and selected for appropriate shRNA expression) were seeded in quadruplicate with 100 μL full serum DMEM media in 96-well round bottom ultra-low attachment spheroid microplate (Corning, 4515). Spheroids were imaged on an IncuCyte S3 with a 10x objective. Spheroid cell viability was quantified by incubation with CellTiter-Glo 2.0 (Promega, PRG9243), and luminescence was read using Tecan Spark plate reader in luminescence mode.

### Colony formation assay

5000 KP4, MiaPaCa-2 or ASPC-1 cells (transduced and selected for appropriate shRNA expression) were seeded with triplicate with 2 mL full serum DMEM media per well of a 6-well plate. Media was changed every 4-5 days over the course of 1.5-2 weeks of growth (until the fastest growing wells were about 80-90% confluent). At the end of the growth period, cells were washed once with PBS and fixed in 4% paraformaldehyde in PBS for 15 min at room temperature. Fixed cells were washed thrice with PBS and subsequently stained with 1 mL of crystal violet staining solution (0.1% crystal violet, 20% methanol in water) for 15 min at room temperature. Stained wells were washed thrice with water and plates were allowed to dry overnight on benchtop. After wells were imaged, remaining crystal violet stain was dissolved in 200-400 μL 10% acetic acid solution. Dissolved crystal violet stain was transferred to clear 96-well plates and absorbance was measured at 595 nm using a Tecan Spark plate reader.

### Continuous cell proliferation assay

6000 KP4, MiaPaCa-2 or ASPC-1 cells (transduced and selected for appropriate shRNA expression) were seeded in 100 μL full serum DMEM media in 96-well clear bottom plate (Corning, 07200588) and cell confluence was monitored using an IncuCyte S3 by taking phase-contrast images with a 10x objective.

### RNA isolation and RT-qPCR analysis

Total RNA extraction was performed using Aurum Total RNA mini kit (Bio-Rad, 7326820) and RNA concentration was estimated from absorbance at 260 nm. cDNA synthesis was performed using iScript Advanced cDNA synthesis kit for RT-qPCR (Bio-Rad, 1708840), following manufacturer’s instructions. cDNA was diluted in DNAse-free water before quantification by RT-PCR. Chemical detection of PCR products was achieved with SYBR Green (Bio-Rad, 172-5271), using a CFX96 touchTM real-time PCR (Bio-Rad). At the end of each run, melt curve analysis was performed. Gene expression was corrected for the expression of reference genes. The following primers were used. For LyLAP – forward: GCAACTGCATACTGGATGCCT; reverse: TCAGGGTTTGAGAGCCATAGC. For Actin – forward: CATGTACGTTGCTATCCAGGC; reverse: CTCCTTAATGTCACGCACGAT.

### SDS-PAGE and immunoblot analysis

Protein lysates were loaded per lane in 4-20% Tris-Glycine gel (Thermo Scientific, XP04205) and resolved by electrophoresis in tris-glycine running buffer (25 mM Tris, 190 mM glycine, 0.1% w/v SDS). Proteins were transferred to a PVDF membrane (Millipore, IPVH00010) and blocked with 5% milk in TBS-T. Membranes were incubated with primary antibody (diluted in 5% milk in TBS-T) overnight at 4°C. Membranes were rinsed in TBS-T for 30 min and incubated with HRP-conjugated secondary antibodies (diluted 1:5000 in 5% milk in TBS-T) for 1 h at room temperature, after which they were washed again for 30 min in TBS-T. Membranes were then incubated with Pierce ECL western blotting substrate (Thermo Scientific, 32109) before being exposed to ProSignal ECL Blotting Film.

### Immunohistochemical analysis

The slides underwent baking at 60°C for 1 h, followed by the removal of paraffin using two 5 min washes in xylene. Rehydration was then carried out sequentially in ethanol (100, 90, 70, 50, and 30%) for 5 min each. Post rehydration, samples underwent a 20 min heat treatment in a 10 mM sodium citrate buffer solution (pH 6.0), followed by two washes with PBS. Subsequently, they were immersed in methanol/5% hydrogen peroxide at room temperature for 30 min. After two PBS washes, the samples were blocked with 2.5% normal goat serum (NGS) for an hour, and then left overnight for incubation with the primary antibody at 4°C. Following this, the slides were washed twice with PBS for 5 min each, before being subjected to secondary antibody incubation. Following three washes, the slides were stained using a di-aminebenzidine (DAB) substrate kit (SK-4100 Vector Laboratories) for 10 min, washed with water, and counterstained with hematoxylin. Finally, the slides were dehydrated, mounted, and bright field images were captured using a KEYENCE BZ-X710 microscope.

### Immunofluorescence and confocal microscopy

Cells were seeded on fibronectin coated glass coverslips in 12-well plates at 250,000 cells per well the day before treatment with compounds for specified concentrations and time (as indicated). Cells were then fixed in 4% paraformaldehyde in PBS for 15 min at room temperature, rinsed thrice with PBS, permeabilized with 0.1% w/v saponin in PBS for 15 min at room temperature, and rinsed again thrice with PBS. Primary antibodies were diluted in 5% normal donkey serum (Jackson ImmunoResearch, 017-000-121) in PBS and placed onto coverslips for 1 h at room temperature. Coverslips were rinsed thrice with PBS and labelled with the appropriate fluorescently conjugated secondary antibodies (diluted 1:300 in 5% normal donkey serum in PBS) for 1 h at room temperature. Coverslips were rinsed thrice with PBS and mounted on glass slides using Vectashield Antifade Mounting Medium with DAPI (Vector Laboratories, H-1200). Confocal microscopy was performed on a spinning-disk Nikon Ti-E inverted microscope (Nikon Instruments) with Andor Zyla-4.5 cMOS camera (Andor Tehcnology) using iQ3 acquisition software (Andor Technology). All images were taken with a 60x oil objective.

Co-localization analysis from immunofluorescence was performed by importing raw, unprocessed, non-overlapping images to FIJI v.2.1.0/1.53c and coverted to 8-bit images. Images of individual channels were thresholded independently to exclude background and converted to binary mask. Co-localization between lysosomes (Lamtor2 or LAMP2-mCherry) and marker of interest (CHMP1A or Phluorin) was determined using the ‘AND’ function of the image calculator. Data plotted as fraction of lysosomes that are positive for marker of interest (‘Colocalization Score’).

Lysosome size was assessed by first projecting 15-20 z-planes per image onto one place using the ‘Image to Stacks’ function followed by the Z-project function on FIJI. The projections were subjected to a pipeline developed in-house on CellProfiler (*71*). Briefly, each projected image was transformed to a grey-scale image and uneven illumination was corrected using the illumination correction function. The resulting image was subjected to ‘IdentifyPrimaryObjects’ function in which the typical diameter of objects (i.e., lysosomes) was manually set and visually inspected for correct identification of lysosomes. The identified primary objects were counted (‘MeasureObjectSizeShape) and area was measured.

For live imaging experiments, an Opera Phenix Automated Microscopy system (Perkin Elmer) was used. 25,000 cells were seeded in duplicate (100 μL/well) of a 96-well glass bottom plate (Cellvis, P96-1.5H-N) in phenol red-free DMEM media. The next day, 50 μL of Hoechst dye was added to cells to stain nuclei 30 min prior to the start the experiment. 4 images of non-overlapping frames were taken per well; first the nuclei were imaged, then cells were imaged at baseline, followed by addition of the appropriate concentration of either LLOMe or GPN (or DMSO vehicle control) and imaging every 1 min for 10 min (for RpH) or 5 min for 30 min (for CHMP4B-mNG). Temperature and CO_2_ levels were maintained at 37°C and 5%, respectively, during the experiment. All images were taken with a 40x water objective. Images were analyzed using Harmony 3D analysis software for nuclei and spot selection and masking and quantification of nuclei/spot number and spot relative intensity for all frames and time-points collected.

### Transmission electron microscopy

PDA cells (transduced with and selected for appropriate shRNAs) were seeded in a 6-well plate. Cells were fixed in 2% glutaraldehyde diluted in warm DMEM media, shaking for 30 min at room temperature. After fixation, cells were scraped into a 1.5 mL microcentrifuge tube and spun down until a pellet was visible. The supernatant was removed, and samples were prepared and sectioned by the University of California Berkeley Electron Microscope Laboratory. Briefly, cell pellets were fixed for 30 min at room temperature in 2.5% glutaraldehyde in 0.1 M cacodylate buffer pH 7.4. Fixed cells were stabilized in 1% very low melting-point agarose and then cut into small cubes. Cubed sample was then rinsed three times at room temperature for 10 min in 0.1 M sodium cacodylate buffer, pH 7.4 and then immersed in 1% osmium tetroxide with 1.6% potassium ferricyanide in 0.1M cacodylate buffer for an hour in the dark on a rocker. Samples were later rinsed three times with cacodylate buffer and then subjected to an ascending series of acetone for 10 minute each (35%, 50%, 75%, 80%, 90%, 100%, 100%). Samples were progressively infiltrated with Epon resin (EMS, Hatfield, PA, USA) while rocking and later polymerized at 60°C for 24 h. 70 nm thin sections were cut using an Ultracut E (Leica, Wetzlar, Germany) and collected on 100 mesh formvar coated copper grids. The grids were further stained for 5 min with 2% aqueous uranyl acetate and 4 min with Reynold’s lead citrate. The sections were imaged using a Tecnai 12 TEM at 120 KV (FEI, Hillsboro, OR, USA) and images were collected using RIO 16 CMOS camera from Gatan. (Gatan Inc., Pleasanton, CA, USA). Lysosome diameter was measured using the measuring tool on ImageJ v2.3.0.

### Lysosomal immunoprecipitation (LysoIP)

Lysosomes from cells expressing TMEM192-mRFP-3xHA were purified as previously described (*37*). Briefly, cells were seeded in 15 cm dishes and, upon reaching appropriate confluence, were harvested into 10 mL ice-cold KPBS buffer (136 mM KCl, 10 mM KH_2_PO_4_, pH 7.25 and supplemented with Pierce Protease Inhibitor tablet). If immunoprecipitated lysosomes were used for further enzymatic assays, no protease inhibitor was added. Cells were pelleted by centrifuging lysates at 300*g* for 5 min and resuspended in a total volume of 1 mL KPBS (supplemented with 5% w/v OptiPrep (Sigma Aldrich, D1556)) and fractionated by passing through a 23G syringe 8 times followed by centrifugation at 800*g* for 10 min (for downstream assays quantifying proteins and lipids), 2 min (for downstream assays quantifying polar metabolites). Post-nuclear supernatant (PNS) was collected, and the rest of the lysate was incubated with 80-100 μL α-HA magnetic beads (Thermo Fisher, 88836) with end-over-end rotation for 30 min (for protein and lipid downstream quantification) and 3.5 min (for polar metabolite quantification) at 4°C. Lysosome-bound beads were rapidly washed three times with KPBS. For immunoblotting, 80-100 μL 2x urea sample buffer was added to the beads and heated for 30 min at 37°C. For proteomic analysis and *in vitro* enzymatic reactions, 80-100 μL KPBS with 1% tergitol was added to the beads and heated for 30 min at 37°C. For lipidomic and metabolomic analysis, 100 μL ice-cold methanol was added directly to the beads and samples were stored at -80°C prior to extraction (details below).

### Recombinant LyLAP expression in HEK293 GnTI-cells

LyLAP (wild-type or C228A mutant) was expressed with a TEV protease cleavable N-terminal 6xHis-tag and IgK secretion signal in HEK293 GnTI-cells upon transient transfection with PEI MAX (Polysciences Inc., 24765). The IgK secretion signal allowed for the secretion of LyLAP recombinant protein. 72 h post transfection, media supernatant was collected by pelleting the cells by centrifugation at 1000 rpm for 20 min at 4°C. Media supernatant was further filtered using 0.2 μm PVDF vacuum filter prior to incubation with Ni-NTA agarose resin (GoldBio, H-350-5) (10 mL wet volume per 1 L supernatant) for 1 h at 4°C. Protein-bound beads were isolated and washed using a gravity-flow column. Beads were washed once with wash buffer 1 (10 mM imidazole, 20 mM HEPES pH 7.5, 200 mM NaCl) and once more with wash buffer 2 (50 mM imidazole, 20 mM HEPES pH 7.5, 200 mM NaCl) prior to elution with 350 mM imidazole (supplemented with 20 mM HEPES pH 7.5 and 200 mM NaCl). Subsequently, the protein elution was run on a Superdex 200 16/600 200 pg column (Cytiva, 28989335). Peak fractions were collected, concentrated using Amicon Ultra Centrifugal filter (Millipore Sigma, UFC903024). Protein was flash-frozen in pH 7.5 buffer or buffer exchanged into “lysosomal pH buffer” (25 mM sodium acetate pH 4.5, 150 mM NaCl) using a Zeba 7k MWCO spin column (Thermo Scientific, PI89882) according to manufacturer instructions and diluted to a final concentration of 1 mg/mL.

### Maleimide labeling and cellular uptake rescue experiments

AF546-Maleimide (Invitrogen, A10258) was reconstituted in DMSO to a 10 mM stock. 70 μg of recombinant LyLAP was incubated with 0.7 μL of AF546-Maleimide (0.1 mM final maleimide concentration). Excess maleimide was removed by buffer exchanging protein into fresh “lysosomal pH buffer” using a Zeba 7k MWCO spin column (Thermo Scientific, PI89882).

For recombinant uptake visualization, KP4 cells were seeded at a density of 50,000 cells/well in 12-well plates and AF546-maleimide-labeled LyLAP was added to cells at a dilution of 1:10 and incubated with cells overnight. The next day, media was changed to fresh DMEM media and cells were fixed for immunofluorescence imaging 3 h later, as normal.

For phenotype rescue experiments (growth, lysosomal size, and membrane permeabilization), unlabeled recombinant LyLAP was added to the media of cells. Recombinant protein was maintained in cell media at a 1:10 concentration during the entire transduction, selection, and experiment phase.

### *In vitro* LyLAP cleavage assay

For autocatalytic cleavage (at pH 4.5), recombinant LyLAP was incubated overnight at 37°C, after which it was aliquoted and flash-frozen in liquid nitrogen for storage at -80°C. Cleavage of recombinant protein was verified by running sample on a Bis-Tris 4-12% gel (Thermo Fisher, NP0321), resolved by electrophoresis in 1x MOPS SDS running buffer (Thermo Fisher, NP000102), and stained using Coomassie (0.02% (w/v) Coomassie G-250 dye, 5% (w/v) aluminum sulfate, 10% (w/v) phosphoric acid, 10% ethanol).

For C228A-LyLAP cleavage by lysate, immunopurified lysosomal lysates were eluted in “lysosomal pH buffer” supplemented with 0.1% tergitol. 30% of the elution was incubated at 37°C with 5 μg of recombinant protein for appropriate time before the sample was collected and run on a denaturing SDS-Page gel and immunoblotted against α-His. For *in vitro* cathepsin cleavage assay, 3 μg of recombinant LyLAP was diluted in “lysosomal pH buffer” containing 0.3-0.7 mU cathepsin (either cathepsin B, D, L, or K) and incubated at 37°C for appropriate time point. Samples were collected and run on Bis-Tris 4-12% gel and visualized using Coomassie. Protease inhibitors (Leupeptin, Pepstatin or E64D) were all added to 20 μM final concentration. Recombinant cathepsins used were as follows: Cathepsin L (Sigma, C6854), Cathepsin K (Abcam, ab157067), Cathepsin B (Sigma, C6286), Cathepsin D (Sigma, C3138).

### Cathepsin B assay

Cathepsin B assay was modified from and conducted as previously described (*72*). Briefly, KP4 cells (transduced and selected for appropriate shRNAs) were seeded at 100,000 cells per well in appropriate replicates in a 12-well plate. Cells were counted and then lysed in 200 μL per well of cold triton lysis buffer (0.1% Triton X-100, 100 mM sucrose, 20 mM HEPES pH 7, 10 mM KCl, 1.5 mM MgCl_2_, 1 mM EDTA), and lysates were nutated at 4°C for 20 min. 50 μL of soluble fraction was incubated with 50 μL of cathepsin B reaction buffer (50 mM sodium acetate, 4 mM EDTA, 8 mM DTT, 50 μM Z-RR-AMC (Sigma, C5429), pH 6) and incubated at 37°C for 5 min. Fluorescence intensity was measured every 30 s for 20 min (ex 350 nm/em 450 nm) using a Tecan Spark plate reader. Lysates treated with protease inhibitor were pre-treated with 20 μM final concentration Leupeptin/Pepstatin A at 37°C for 30 min prior to addition of the Z-RR-AMC substrate.

### Acid ceramidase assay

Fluorogenic acid ceramidase assay was modified from one previously described (*73*). Briefly, KP4 cells (transduced and selected for appropriate shRNAs) were seeded in 10 cm dishes. Cells were counted then resuspended in 1 mL of cold 0.2 M sucrose solution and lysed by sonication in a water bath for 20 min followed by freeze-thawing in liquid nitrogen. In a well of a black 96-well plate, 25 μL of each lysate was added to 74.5 μL of “lysosome pH buffer”, 0.5 μL of 4 mM RBM14C12 substrate (Avanti Lipids, 860855) (resuspended in ethanol) for a final concentration of 20 μM. Where indicated, samples treated with 0.1 μL of 3 mM Carmofur (Sigma-Aldrich, C1494). Samples were incubated at 37°C for 3 h (without agitation and in dark). To stop the enzymatic reaction, 50 μL methanol and 100 μL of 2.5 mg/mL NaIO_4_ in 100 mM glycine/NaOH pH 10.6 was added to each well. Fluorescence (ex 360 nm/em 446 nm) was detected at room temperature every 15 min for 2 h (until stabilized) using Tecan Spark plate reader.

### β-galactosidase assay

KP4 cells (transduced and selected for appropriate shRNAs) were seeded at 10,000 cells per well in a flat-bottom black 96-well plate with clear bottom (Corning, 07200588) and allowed to adhere overnight. Media was aspirated and replaced with 100 μL fresh full serum DMEM media supplemented with 15 μM LysoLive GalGreen fluorogenic substrate (MarkerGene Technologies, M2776) and 50 nM LysoTracker Red DND-99 (Thermo Scientific, L7528) and incubated at 37°C for 2 h. Wells were rinsed with PBS and media was replaced with imaging buffer (136 mM NaCl, 2.5 mM KCl, 2 mM CaCl_2_, 1.3 mM MgCl_2_, 10 mM HEPES pH 7.5). Fluorescnece was measured on a Tecan Spark plate reader (ex 485 nm/em 528 nm and ex 570 nm/em 590 nm) from the bottom of the plate. GalGreen fluorescence was normalized per well to LysoTracker red fluorescence.

### Bile salt hydrolase activity

Bile acid hydrolysis assay was modified from method previously described (*74*). Briefly, 80 pmol of recombinant LyLAP (supplemented with 0.2 mU cathepsin L) was diluted in 50 μL final volume of “lysosomal pH buffer” and added to a well of a black 96-well plate. To begin the reaction, 50 μL of cholic acid-AMCA probe (diluted in “lysosomal pH buffer”) was added to each well for a final probe concentration of 150 μM. 2 units (per reaction) of bile salt hydrolase (BSH; Sigma-Aldrich, C4018) was used as a positive control. Fluorescence (ex 350 nm/em 450 nm) was measured every minute for 20 min (until stabilized) using Tecan Spark plate reader.

### LyLAP monoaminopeptidase activity

*In vitro* LyLAP activity assays were done in a 384-well format with a Tecan Spark plate reader. LyLAP (200 nM for single concentration turnover assay or 20 nM for steady-state activity assays) was preactivated, where indicated, with Cathepsin L (0.2 mU) for 15 min at 37°C in “lysosomal pH buffer”. Activated LyLAP (15 μL, 20 nM final) was combined with 2x substrate stock solution (15 μL; diluted in “lysosomal pH buffer”) with 4-5 replicates. For Leu-AMC (L-Leucine-7-amido-4-methylcoumarin; Sigma-Aldrich, L2145), 11 concentrations ranging from 125 μM to 2.2 μM were tested. For Ile-AMC (L-Isoleucine-7-amido-4-methylcoumarin; SCBT, sc-295292C), 7 concentrations ranging from 80 μM to 0.5 μM were tested. Hydrolysis was monitored continuously at 37°C for 1 hour (ex 340 nm/em 440 nm). The initial velocities (RFU/s) from 5-15 minutes were converted to molar units (M/s) via a standard curve of free AMC (Millipore Sigma, 257370) and free Leu-AMC/Ile-AMC. The kinetic parameters k_cat_ and K_M_ were determined from the plot of initial velocity (M/s) against substrate concentration (M) using a Michaelis-Menton fit in GraphPad Prism. For single concentration turnover assays, LyLAP (15 uL, 200 nM final) was combined with 2X substrate stock solutions (15 uL) with three replicates. Negative control substrates were Pro-AMC (L-Proline-7-amido-4-methylcoumarin; Sigma-Aldrich, P5898) and Gly-AMC (Thermo Scientific Chemicals, J65592.MA). Kinetic parameters for tripeptides LLL and LPL were obtained in the same way as described above using tripeptide-AMC conjugates (synthesized by GenScript). All other processing steps are as described above.

### Lipid hydrolysis assays by LC-MS

For lipid hydrolysis assays, 20 μg recombinant LyLAP (pre-activated with 0.2 mU cathepsin L for 15 min at 37°C) was diluted in 100 μL total volume either “lysosomal pH buffer” (pH 4.5) or PBS in the presence of 5% (final concentration) ethanol. For phospholipase B assay, 16:0/18:1 PC (POPC; Avanti Lipids, 850457) was added to the reaction mixture at 100 μM final concentration and incubated at 37°C for 20 h. For ceramidase assay, C12 Ceramide (Avanti Lipids, 860512) was added to the reaction mixture at 200 μM final concentration and incubated at 37°C for 0, 0.5 h, 1 h, 4 h, and 20 h. For glucosylceramide N-deacylase assay, C12 Glucosylceramide (Avanti Lipids, 860543) was added to the reaction mixture at 100 μM final concentration and incubated at 37°C for 20 h. The positive control for the ceramidase assay was recombinant ASAH1 (purified in-house using the same purification strategy and autocatalytic activation strategy as for LyLAP). The positive control for the glucosylceramide N-deacylase assay was recombinant Sphingolipid ceramide N-deacylase (SCDase; Sigma-Aldrich, S2563).

Each reaction was stopped by adding 400 μL of tert-butyl methyl ether (MTBE; Sigma-Aldrich, 34875) and 120 μL of methanol, supplemented with C12-mono-acyl-glycerin-ether (C12 MAGE) internal standard for 20 μM final concentration. Apolar phase (top) was transferred to a new tube, dried down by speed-vac, resuspended in 120 μL MTBE and transferred to a glass vial with a glass vial insert inside prior to injection on LC-MS.

10 μL of each sample was injected and analyzed by single-reaction monitoring (SRM)-based LC-MS/MS (*75*). Metabolite separation was achieved with a Luna reverse-phase C5 column (50 x 4.6 mm, with 5-μm-diameter particles; Phenomenex). Mobile phase A was composed of 95:5 water:methanol; mobile phase B was composed of 60:35:5 isopropanol:methanol:water. Solvent modifiers 0.1% formic acid with 5 mM ammonium formate were used to assist in ion formation and improve LC resolution in positive mode. The flow rate for each run started at 0.1 mL/min for 5 min. The gradient started at 0% B and increased linearly to 100% B over 45 min with a flow rate of 0.4 mL/min, followed by isocratic gradient of 100% B for 17 min at 0.5 mL/min, before equilibrating for 8 min at 0% B with the same flow rate (0.5 mL/min). MS analysis was performed with electrospray ionization source on an Agilent 6430 QQQ LC-MS/MS. Capilary voltage was set to 3.0 kV, drying gas flow rate was 10 L/min and nebulizer pressure was 35 psi. Representative lipids were quantified by SRM of the transition from precursor to product ions at optimized collision energies.

### Sphingolipid hydrolysis assays by thin-layer chromatography (TLC)

For complex sphingolipid hydrolysis assays by TLC, 0.3 nmol recombinant LyLAP (pre-activated with 0.2 mU cathepsin L for 15 min at 37°C) was diluted in 20 μL total volume either “lysosomal pH buffer” (pH 4.5) containing 0.2% Triton X-100 and 10 nmol of appropriate glycosphingolipid. The following glycosphingolipids were used: C12 Galactosylceramide (Avanti Lipids, 860544), Ganglioside GM3 (Avanti Lipids, 860058), Ganglioside GD3 (Avanti Lipids, 860060), and a mixture of gangliosides consisting of Asialo-GM1, GM1, GD1a, GD1b, GT1b (Cayman Chemical, 29362). The positive control for the glycolipid N-deacylase assay was recombinant Sphingolipid ceramide N-deacylase (SCDase; Sigma-Aldrich, S2563) (*76*). Reaction mixtures were incubated at 37°C for 20 h after which the reaction was stopped and lipids were extracted by addition of 375 μL methanol, 1250 μL MTBE and 315 μL water to induce phase separation. Apolar phase (top) was collected and dried down using speed-vac. Entire dried extract was resuspended and spotted onto HPTLC Silica Gel plate (Millipore Sigma, M1137480001). TLC running solvent comprised of 60:35:4 chloroform:methanol:water. Plates were developed in either 0.1 mg/mL fluorescamine (Sigma-Aldrich, F9015) in acetone to visualize primary amine-containing lipids or 0.05% primuline in acetone to visualize all other lipids.

### Deglycosylation assay

15 μg of recombinant LyLAP was deglycosylated using PNGase F (NEB, P0704) in either native or denaturing conditions, following manufacturer’s instructions. Deglycosylated protein was resolved by electrophoresis on a Bis-Tris 4-12% gel in 1x MOPS SDS running buffer and stained using Coomassie.

### Matrix assisted laser desorption ionization-time of flight (MALDI-TOF) spectrometry

Recombinant LyLAP was diluted to 10 μM final concentration in 20 μL total of 5 mg/mL Sinapinic acid matrix solution (Supelco, 49508) prepared in 70:30 acetonitrile:water + 0.1% trifluoroacetate. 0.8 μL of protein-matrix solution was spotted onto a MALDI plate (JBI Scientific, V700666) and allowed to air-dry. Mass spectra was acquired on a Voyager DE Pro (Applied Biosciences) in linear mode with the following settings: 25000 V accelerating voltage; positive polarity; 10000-80000 Da acquisition mass range; 500 laser shots per spectrum.

### Untargeted lipidomics

#### Sample preparation and liquid chromatography

LysoIP samples were diluted up to 300 μL in ice-cold LC-MS grade methanol (Fisher Scientific, A452-4). Internal standards – 5 μl SPLASH® LIPIDOMIX® (Avanti Polar Lipids, 330707) dissolved in methanol was added to each sample. Lipids were extracted by adding an additional 1250 μl tert-butyl methyl ether (Sigma-Aldrich, 34875) and 75 µL methanol to each sample. The mixture was incubated on a revolving mixer for 1 h (room temperature, 32 rpm). To induce phase separation, 375 µL H_2_O (LC-MS grade; Fisher Chemical, W6-4) was added, and the mixture was incubated on a revolving mixer for 10 min more (room temperature, 32 rpm). Samples were centrifuged (room temperature, 10 min, 1000*g*). Upper organic phase with collected and subsequently dried *in vacuo* (Eppendorf concentrator 5301, 1 ppm).

Dried lipid extracts were reconstituted in chloroform/methanol (50 µl, 2:1, v/v) and 10 µl of each extract was transferred to HPLC vials containing glass inserts. Quality control (QC) samples were generated by mixing equal volumes of each lipid extract. QC samples were aliquoted to 10 µL, dried *in vacuo*, and redissolved in 2-propanol (10 µl) for injection. Lipids were separated by reversed phase liquid chromatography on a Vanquish Core (Thermo Fisher Scientific, Bremen, Germany) equipped with an Accucore C30 column (150 x 2.1 mm; 2.6 µm, 150 Å, Thermo Fisher Scientific, Bremen, Germany). Lipids were separated by gradient elution with solvent A (MeCN/H_2_O, 1:1, v/v) and B (i-PrOH/MeCN/H_2_O, 85:10:5, v/v) both containing 5 mM NH_4_HCO_2_ and 0.1% (v/v) formic acid. Separation was performed at 50°C with a flow rate of 0.3 mL/min using the following gradient: 0-15 min – 25 to 86 % B (curve 5), 15-21 min – 86 to 100 % B (curve 5), 21-34.5 min – 100 % B isocratic, 34.5-34.6 min – 100 to 25 % B (curve 5), followed by 8 min re-equilibration at 25 % B.

#### Mass spectrometry

Reversed phase liquid chromatography was coupled on-line to a Q-Exactive Plus Hybrid Quadrupole Orbitrap mass spectrometer (Thermo Fisher Scientific, Bremen, Germany) equipped with a HESI probe. Mass spectra were acquired in positive and negative modes with the following ESI parameters: sheath gas – 40 L/min, auxiliary gas – 10 L/min, sweep gas – 1 L/min, spray voltage – 3.5 kV (positive ion mode); -2.5 kV (negative ion mode), capillary temperature – 250°C, S-lens RF level – 35 and aux gas heater temperature – 370°C.

Data acquisition for lipid identification was performed in quality control samples by acquiring data in data-dependent acquisition mode (DDA). DDA parameters featured a survey scan resolution of 140,000 (at *m/z* 200), AGC target 1e6 Maximum injection time 100 ms in a scan range of *m/z* 240-1200. Data dependent MS/MS scans were acquired with a resolution of 17,500, AGC target 1e5, Maximum injection time 60 ms, loop count 15, isolation window 1.2 *m/z* and stepped normalized collision energies of 10, 20 and 30 %. A data dependent MS/MS scan was triggered when an AGC target of 2e2 was reached followed by a Dynamic Exclusion for 10 s. All isotopes and charge states > 1 were excluded. All data was acquired in profile mode.

For deep lipidome profiling, iterative exclusion was performed using the IE omics R package (*77*). This package generates a list for fragmented precursors from a prior DDA run that can be excluded from subsequent DDA runs ensuring a higher number of unique MS/MS spectra for deep lipidome profiling. After the initial DDA analysis of a quality control sample, another quality control sample was measured but excluding all previously fragmentated precursor ions. Parameters for generating exclusion lists from previous runs were – RT window = 0.3; noiseCount = 15; MZWindow = 0.02 and MaxRT = 36 min. This workflow was performed twice to achieve a total of three DDA analyses of a quality control sample in positive and three DDA analyses in negative ionization mode.

Data for lipid quantification was acquired in *Full MS* mode with following parameters – scan resolution of 140,000 (at *m/z* 200), AGC target 1e6 Maximum injection time 100 ms in a scan range of *m/z* 240-1200.

#### Lipid identification and quantification

Lipostar (version 2.0, Molecular Discovery, Hertfordshire, UK) equipped with *in-house* generated structure database featuring fatty acids with no information on double bond regio-or stereoisomerism covering glycerolipid, glycerophospholipid, sphingolipid and sterol ester lipid classes. The raw files were imported directly with a *Sample MS Signal Filter Signal Threshold* = 1000 for MS and a *Sample MS/MS Signal Filter Signal Threshold* = 10. Automatic peak picking was performed with an *m/z tolerance* = 5 ppm, *chromatography filtering threshold* = 0.97, *MS filtering threshold* = 0.97, *Signal filtering threshold* = 0. Peaks smoothing was performed using the *Savitzky-Golay* smoothing algorithm with a *window size* = 3, *degree* = 2 and *multi-pass iterations* = 3. Isotopes were clustered using a *m/z tolerance* = 5 ppm, *RT tolerance =* 0.25 min, *abundance Dev* = 40%, *max charge* = 1. Peak alignment between samples using an *m/z tolerance* = 5 ppm and an *RT tolerance* = 0.25 min. A gap filler with an *RT tolerance* = 0.05 min and a *signal filtering threshold* = 0 with an anti-Spike filter was applied. For lipid identification, a “MS/MS only” filter was applied to keep only features with MS/MS spectra for identification. Triacylgylcerols, diacylglycerols and sterol esters were identified as [M+NH4]+ adducts. Lysophosphatidylcholines, lysophosphatidylethanolamines, Ether-and vinyl ether-PE, ceramides and Sphingomyelins were analyzed as [M+H]+ adducts. Phosphatidylserines, phosphatidylinositols were analyzed as [M-H]-adducts. Acyl-, ether-and vinyl ether-Phosphatidylcholines were identified as [M+HCOO]-adducts. Following parameters were used for lipid identification: 5 ppm precursor ion mass tolerance and 20 ppm product ion mass tolerance. Automatic approval was performed to keep structures with quality of 3-4 stars. Identifications were refined using manual curation and Kendrick mass defect analysis and lipids that were not following these retention time rules were excluded as false positives.

Quantification was performed by peak integration of the extracted ion chromatograms of single lipid adducts. Peak integration was manually curated and adjusted. Identified lipids were normalized to peak areas of added internal standards to decrease analytical variation and normalized to protein concentrations of lyso-IP samples, as measured by BCA assay (Thermo Fisher, 23225). Statistical analyses for lipidomics data was conducted using MetaboAnalyst webtool (metaboanalyst.ca) (*78*).

### Untargeted metabolomics

Whole-cell metabolomic extraction was performed by flash freezing 6-well culture plates in liquid nitrogen after one rapid wash with ice-cold water. On dry ice, 1 mL of a -20°C 1:1 methanol:water solution was added. Cells were quickly scraped on wet ice, transferred to a 2mL tube containing 1 mL of ice-cold chloroform, vortexed and kept on ice.

Endogenous proteins in LysoIP samples were heat denatured at 95°C for 5 min. Lysates were then treated overnight at 37°C with 150 nM (final concentration) recombinant LyLAP and 0.3 mU cathepsin L (or remained untreated) and were subsequently quenched with 100 µL ice-cold methanol and dried down. Dried extracts were reconstituted with 400 μL 90% methanol solution. 320 µL of ice-cold water was added to achieve a 1:1 water:methanol ratio, and this solution was transferred to a tube with 720 µL of ice-cold chloroform and vortexed.

Both LysoIP samples and whole cell samples were vortexed in 3 cycles of 20 sec, each followed by a 5 min resting period on ice. Phase separation was completed by centrifugation at 25000*g* (4°C, 20 min). The polar phase (top) was carefully pipetted to a new tube and dried in an unheated vacuum concentration system. Metabolites were resuspended in 50 µL of a 50% methanol solution, centrifuged at 25000*g* (4°C, 10 min), and 5 µL of the supernatant was taken for LC-MS analysis. The chromatographic separation of metabolites was performed using the hexylamine-based method as previously described (*79*). Mass spectrometry data was acquired by an Agilent 6550 Q-TOF as described before (*53*).

For TMT derivatization of small peptides for MS2 identification, dried lysosomal metabolite extracts were resuspended in 50 µL of 50 mM TEAB buffer (pH 8.5). Subsequently, 40 µL of acetonitrile was added to each sample. TMTzero Label reagent (Pierce) was dissolved in anhydrous acetonitrile at 20 µg/µL, and 10 µL of TMTzero solution was added to each sample. After 1 h incubation at room temperature, the derivatization reaction was quenched by addition of 4 µL of a 2.5% hydroxylamine solution and incubation for 15 min at room temperature. Samples were dried down once more and resuspended in 50 µL of 50% methanol for LC-MS analysis using the metabolomic LC-MS setup described above. For the MS2 identification of small TMT-labeled peptides, selected precursors were isolated in a 4 mass unit window by the quadrupole, followed by collision-induced dissociation (CID) at 20 kEV.

For untargeted metabolomic analysis of lysosomal and whole-cell metabolite extracts, peak features were detected, grouped, and quantified using the Profinder software (Agilent). Peak abundances were log-transformed and subsequently normalized by subtracting the median abundance within each sample. Volcano plots were made to visualize metabolite enrichment in LyLAP knockdown samples using the Perseus software (Max Planck Institute) (*80*) and R. The p-values making up the y-axis of the volcano plot were calculated with an unpaired two-tailed t-test. Target metabolites were identified by matching m/z and retention times with standards. Quantification was based on the extracted ion chromatogram (EIC) area of the [M-H]^-^ ion. Within each sample, abundances were normalized to average of all measured metabolites.

### Multiplexed Substrate Profiling-Mass Spectrometry (MSP-MS)

The MSP-MS library was used to determine substrate specificity of LyLAP as reported previously (*81*). 200 nM (final concentration) LyLAP was preactivated with 0.3 mU Cathepsin L for 30 minutes in NaOAc (Sigma Aldrich) pH 4.0. To a preheated buffer containing NaOAc pH 4.0, a final of 20 uM of Leupeptin (Thomas Scientific), 1 µM of the peptide library, and 20 nM of LyLAP^active^ was added. The reaction was then allowed to proceed at 37°C and 15 µL aliquots were taken out as timepoints at 15 min, 1 h, 4 h, and 24 h. These were then quenched with 15 µL of 6 M Guanidinium chloride. The samples were then acidified by spiking in Formic acid to a final concentration of 2%. These samples were desalted using preequilibrated (3x 15 µL of 50% ACN 0.2% Formic acid then 3X 15 µL 0.2% formic acid) Cleanup C18 pipette tips (Agilent) by pipetting 15 µL of the acidified solution 10X to ensure full binding of the peptides to the C18 plug in the tips. The tips were then washed 5X with 15 µL of 0.2% formic acid, and finally with 5 x 15 µL of 50% ACN, 0.2% Formic acid eluted into a to a nonstick 0.5 ml Axygen maximum recovery tube (Corning). This eluant was then evaporated in a speed vac and reconstituted with 15 µL of 0.1% formic acid. 5 µL of each sample was then injected into a PepMap RSLC C18 (Thermo Scientific ES900) attached to a 10,000-psi nanoACQUITY Ultra Performance Liquid Chromatography System (Waters) followed by a Q Exactive Plus Hybrid Quadrupole-Orbitrap (Thermo Fisher Scientific). The peaks were assigned using PAVA and the peaks were searched using ProteinProspector to the MSP-MS database and then analyzed using Extractor as described previously (*81*). and visualized using the ICE-logo web tool (*82*) with a reference set of all possible P4-P4’ cleavages in the MSP-MS library. The Ice-logo was created by weighting the spectral counts observed across all replicate experiments.

### Hydrophobic peptide processing and peptide library synthesis

A small sub-library was generated based on the sequence of a peptide from the MSP-MS library (FLKAHSDVWPYQDA) as it was already demonstrated to be a substrate for LyLAP in the MSP-MS experiment. F and L were replaced by the top five hydrophobic amino acids from MSP-MS were added to cover all combinations with an extra L was added to the N-terminus of each of the 25 peptides to observe sequential cleavage.

The peptides were synthesized on the Syro II peptide synthesizer (Biotage) using the standard Fmoc solid phase synthesis. All the peptides were synthesized using 12 µmol of preloaded alanine resin (Sigma-Aldrich) at ambient temperature. All couplings were done with 5 equivalence of amino acid (Novabiochem), 4.9 equivalences of O-(1H-6-Chlorobenzotriazole-1-yl)-1,1,3,3-tetramethyluronium hexafluorophosphate (HCTU) (Anaspec), and 20 equivalence of N-Methylmorpholine (NMM) (Sigma-Aldrich) in 500 µL of Dimethylformamide (DMF) (Sigma-Aldrich). Each of the amino acids were double coupled using 8 min reactions. The Fmoc was deprotected using 500 µL of 40% 4-Methylpiperidine (TCI) in DMF for 3 min, then a wash of 500 µL of 20% of 4-Methylpiperidine in DMF for 10 min. The deprotected peptides on resin were then washed 6 times with 500 µL of DMF for 6 min. The peptides were cleaved from the resin using a cleavage solution (88% TFA (Sigma-Aldrich), 5% phenol (Sigma-Aldrich), 5% Triisopropylsilane (Sigma-Aldrich), 2% H2O (MilliQ) and were shaken for 3 h before precipitation with 45 mL of ice cold 1:1 diethyl ether (Sigma-Aldrich) and hexanes (Sigma-Aldrich). The suspension was pelleted through centrifugation at 4,000 xg for 10 min and the supernatant was removed. The resulting pellet was allowed to dry in a chemical hood overnight. The crude pellet was re-dissolved in 200 µL of dimethyl sulfoxide (DMSO) (Sigma-Aldrich). Each peptide was purified by HPLC on an Agilent Pursuit 5 C18 column (5 mm bead size, 150 x 21.2 mm) attached to an Agilent PrepStar 218 HPLC. Peptides were eluted using a mobile phase with an ACN (Sigma-Aldrich) 0.1 % TFA gradient 20% to 80%. The solvent was removed from the purified material through evaporation using the Genevac EZ-Bio Standard. The purified peptides were resuspended in 3 mL of water, confirmed by LC-MS, and lyophilized. With the generated sub-library, the peptides were constituted into 50 mM DMSO and were mixed to a final dilution of 1 µM of each peptide into the same reaction buffer as MSP-MS. Aliquots were taken at six timepoints (0 min, 15 min, 1 h, 2 h, 4 h, 24 h) and quenched in equal volume of 6M Guanidium HCL.

The peptides were prepared for MS analysis as the MSP-MS library vide supra. The .raw data was exported from the instrument and imported directly into the MSFragger/Fragpipe program (*83–88*). The data was analyzed using the LFQ-MB workflow. The spectra were searched against a database that was generated from a FASTA file of all 25 peptides. Decoys were added to this database with the Fragpipe program including common contaminant proteins. All counts from contaminants were removed from downstream analysis. The spectral count for these peptides were then plotted to determine processivity for each generated peptide. Because Leu and Ile are the same weight, these residues are indistinguishable in this assay and are all plotted as Leu

### Semi-tryptic peptide analysis

High confidence LyLAP substrate peptides were probed by immunoprecipitating lysosomes from shLyLAP KP4 cells and eluting in “lysosomal pH buffer” supplemented with 0.1% tergitol. Endogenous proteins were heat denatured at 95°C for 5 min and lysates were treated overnight at 37°C with 150 nM (final concentration) recombinant LyLAP. Prior to addition to the lysosomal lysate, recombinant LyLAP was cleaved with 0.3 mU of cathepsin L at 37°C for 45 min, after which the samples were mixed with a mixture of leupeptin and pepstatin A (final concentration of both 20 µM). After overnight treatment with recombinant LyLAP, the reaction was stopped by again heat denaturing all the proteins at 95°C for 5 min, after which the samples were flash-frozen in liquid nitrogen and stored at -80°C.

BCA was performed to quantify protein concentrations across samples so they could be diluted to contain the same protein concentration. To 100 µL of each sample, an equal volume (100µL) of 2X denaturing buffer consisting of 10% SDS, 100mM Tris at pH 8.5 and 10 mM DTT was added. Samples were denatured, loaded, digested and purified with the micro S-Trap column (ProtiFi) according to the manufacturer’s protocol. Samples were digested with 1:50 LysC (Wako Chemicals) and 1:50 Trypsin (Promega) relative to protein amount. Following a quantitative peptide assay (Pierce) of the reconstituted samples, 600 ng of peptides were injected onto a C18 nanoLC column (EasySpray, 0.075 x 250 mm, Thermo) with a linear gradient of 4%-27.5% solvent B (0.1% FA in 80% ACN) for 37 min, 27.5%-50% solvent B for 20 min, 50%-95% solvent B for 10 min at a constant flow rate of 300 nL/min on a Vanquish Neo HPLC system. The peptides were analyzed by an Orbitrap Fusion Lumos mass spectrometer (Thermo). A semitrypic peptide search was performed using the Sequest HT algorithm in Proteome Discoverer 2.5 (Thermo) to identify peptides based on the human proteome database. For this search, we used a precursor match tolerance of 20 ppm, a fragment mass tolerance of 0.05 Da and a False Discovery Rate (FDR) of 1%. Using the Perseus software (*80*), missing values were imputed based on the Gaussian distribution of log-transformed abundance values. The down shift for the imputed values was set to 2 SD.

For post-processing, peptides that were detected in less than 2 of the replicates in the LyLAP^dead^ samples and those marked as contaminants were removed from the analysis. Only N-semitryptic peptides were then processed and log_2_-fold changes and p-values were calculated. Peptides with a p-value of less than 0.1 were removed. Out of 13,642 starting peptides, 90 were considered as substrates for LyLAP^active^ (as they were enriched in the LyLAP^dead^ samples and achieved the rest of the filtering criteria). Hydrophobicity index of individual peptides was calculated using the Peptides package in RStudio and the hydrophobicity function using the Abraham-Leo scale (*89*).

### Bioinformatic analysis

Expression-survival correlation analysis was conducted by first making a list of 179 proteins that fell under vacuole GO term (GO:0005773) and hydrolase GO term (GO:0016787). Using pathology data from the Human Protein Atlas and The Cancer Genome Atlas (TCGA), only 68 proteins that had listed an unfavorable prognostic outcome were kept (*90–92*). Comparative analysis of lysosomal proteomics data from HPDE versus 8988T cells was based on method previous described (*37*). Briefly, the minimum peptide abundance was set to 1 for all replicates. Only proteins with an average peptide abundance of >1.5-fold enrichment over blank samples were included in the analysis. Fold changes between experimental samples were calculated and significance was calculated using two-tailed unpaired t-tests. Significantly enriched proteins from this analysis refer to those that have a p-value < 0.05 and a log_2_-fold-change > 2 in 8988T lysosomes over HPDE. Venn diagram comparing expression-survival analysis and proteins enriched in PDA lysosomal proteomics data was generated using the VennDiagram package in RStudio. Network analysis of 16 hydrolases that were both unfavorable prognostically in cancer and enriched in PDA lysosomes was conducted by finding the associated pathway in KEGG for each protein and generating a network using the igraph package (igraph.org) in RStudio.

Gene ontology analysis of most enriched proteins from lysosomal proteomics of 8988T versus HPDE cells was performed using Panther overrepresentation test; p-values are calculated by Fisher’s exact test with false discovery rate calculated. Homo sapiens gene whole-genome list was used as reference (*93*, *94*). LyLAP mRNA levels in patient PDA tumors versus surrounding tissues were determined by analyzing data from published and deposited studies in the Gene Expression Omnibus (*95–97*). mRNA levels of 16 top hydrolases (including LyLAP) in PDA versus non-PDA cancer cell lines was assessed by analyzing transcriptional expression data from the Cancer Cell Line Encyclopedia (CCLE; (*98*)). Log_2_ mean expression for each protein within each tumor type was calculated and expression was normalized per column (i.e., per protein) of the heatmap.

Comparative analysis of lysosomal proteomics data from LyLAP-depleted KP4 cells was conducted in the same way as described above. Proteins belonging to general autophagic cargo category were determined by identifying proteins that were significantly accumulated (log_2_-fold-change > 2) upon BafA1 treatment compared to DMSO in shLuc cells (log_2_(shLuc_BafA1_/shLuc_DMSO_)). To identify proteins that were likely to be LyLAP substrate proteins (i.e., more enriched than normal lysosomal autophagic cargo), the following formula was employed: 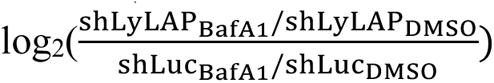. Max hydrophobicity of peptide stretches belonging to each group of enriched proteins was conducted by picking the maximum hydrophobicity value from the ‘membpos’ function within the Peptides package in RStudio (*55*, *56*); maximum hydrophobicity was plotted as a density plot using the ggplot2 package in RStudio. Annotations regarding subcellular localization and transmembrane domain for proteins within each group was gathered using the ‘ID mapping’ function in Uniprot (uniprot.org). Network plot of subcellular localization for each protein was generated using Cytoscape (*99*).

All heatmaps were generated using tidyHeatmap package in RStudio.

### Statistical analysis

Data were expressed as mean ± standard deviation. Statistical analyses were performed using unpaired two-tailed Student’s t-tests for comparisons of 2 groups, one-way ANOVA or two-way ANOVA for group comparisons using GraphPad Prism v10.2.0 (GraphPad Software). Details of each statistical test are given in the legend accompanying each figure. Curve-fitting for enzyme kinetics and other statistical analyses for metabolomics or proteomics are detailed in the appropriate methods.

## Supplementary Text

### Determining LyLAP cleavage pattern after autocatalytic cleavage

The existence of the L209-H227 loop in autocatalytically-cleaved LyLAP (WT (a.c.)) was determined using a combination of glycosylation pattern-matching after PNGase F treatment, exact mass by MALDI-TOF, and previously published data on glycosylation sites on LyLAP (*43*). First, it was noted that the N-terminus of WT (a.c.) had at least two glycosylation sites based on the PNGase F treatment by comparing the stepwise size shift in the native deglycosylated protein and the denatured deglycosylated protein with the glycosylated (blue box) (Fig. S4H). Next, the exact mass of the N-terminus of the WT (a.c.) protein was determined by MALDI-TOF (Fig. S4J) to be the same as the calculated cumulative mass of the 6x-His + TEV + N-terminal domain (including the L209-H227 loop) + two glycans (Fig. S4I). Therefore, the existence of at least two glycans, one of which must come from the fully matured LyLAP N-terminal domain and the other from within the loop, and the exact mass matching suggested that the autocatalytically activated protein still contained the L209-H227 loop on the N-terminal domain of the protein.

### N-semitryptic peptidomic analysis for the identification of endogenous LyLAP peptide substrates

Upon tryptic digestion of lysosomal lysates to allow for peptide profiling, depletion of LyLAP monoaminopeptidase activity results in accumulation of peptides with a single tryptic site on the C-terminus, and an uncut LyLAP site at the N-terminus (referred to as N-semitryptic). Adding back LyLAP^active^ is expected to degrade the N-termini of these peptides, whereas LyLAP^dead^ should not (Fig. 4A, S8A). When we profiled the peptides that followed this pattern, the majority began with one or more hydrophobic N-terminal residues, with leucine being the most common (Fig. 4B, S8B).

---

**Fig. S1.**
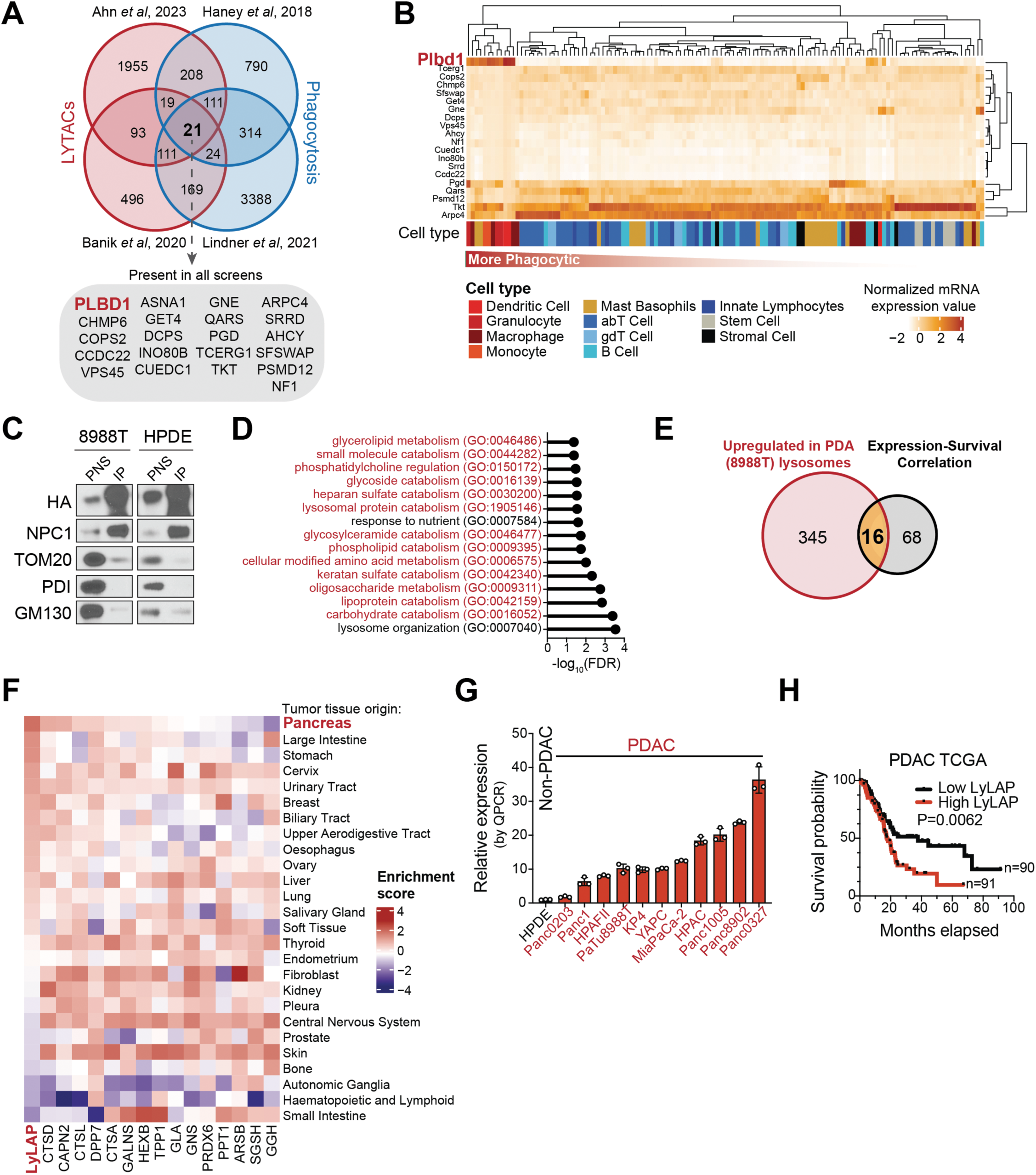
LyLAP is a lysosomal hydrolase enriched in highly endocytic cells. **A.** Venn diagram of gene hits from functional genomics screens identifying regulators of LyTACs-dependent endocytosis and phagocytosis regulators identifies 21 genes that are hits in all screens. **B.** Heatmap of normalized mRNA expression of 21 genes identified in **A.** over immune cell populations (data sourced from Immunological Genome Project). **C.** Immunoblot of Lyso-IP sample, and corresponding post-nuclear supernatant (PNS) used for quantitative lysosomal proteomics analysis (described in Fig. 1A). **D.** GO enrichment analysis of 345 proteins that are significantly upregulated in 8988T lysosomes compared to HPDE lysosomes. GO terms for lysosomal hydrolases annotated in red. **E.** Venn diagram of the overlapping proteins between the lysosomal proteomics analysis and cancer-associated patient survival analysis (described in Fig. 1A). **F.** Enrichment score for RNA-seq data from the Cancer Cell Line Encyclopedia (CCLE) of 16 up-regulated hydrolases (from **F.**) across cancer types. **G.** Quantitative PCR (qPCR) of LyLAP mRNA expression in PDA cell lines (red) compared to control HPDE cell line (black). **H.** Kaplan-Meier survival curve of PDA patients stratified by LyLAP mRNA expression (data from The Cancer Genome Atlas). Data are shown as mean ± SD. Comparison of groups carried out by simple survival analysis (**I**).

**Fig. S2.**
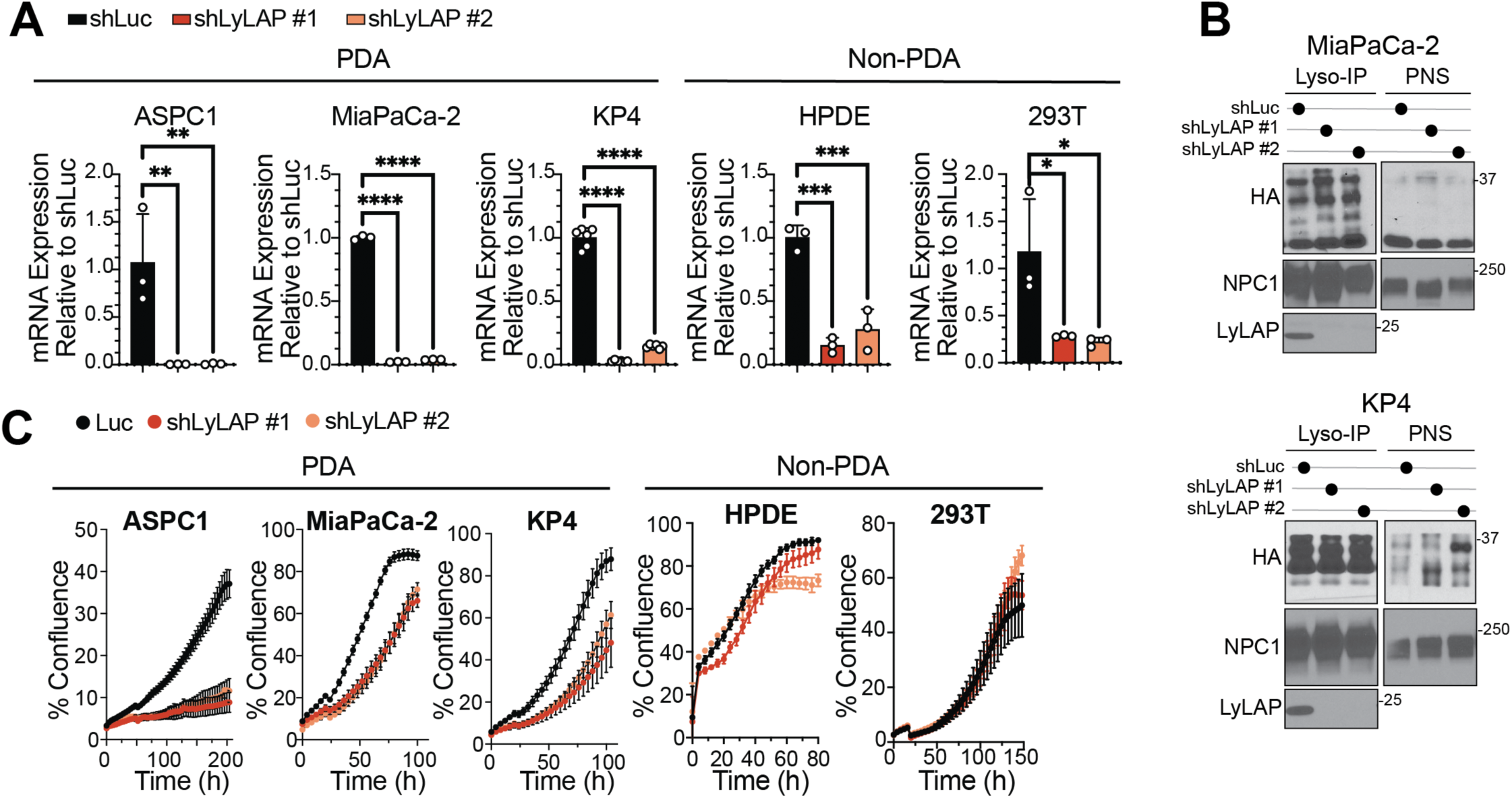
LyLAP is required for PDA growth. **A.** qPCR of LyLAP mRNA expression in PDA (ASPC1, MiaPaCa-2 and KP4) and non-PDA (HPDE, 293T) cell lines upon shRNA-mediated LyLAP knock down compared to shLuc control. **B.** Immunoblot analysis of LyLAP expression in lysosomal immunoprecipitates from MiaPaCa-2 and KP4 PDA cells upon LyLAP-knockdown by shRNA compared to shLuc control. PNS not shown for LyLAP, as protein is not detectable by antibody in whole-cell lysates. **C.** 2D cell proliferation assay for PDA cell lines (ASPC1, MiaPaCa-2, and KP4) treated with the indicated shRNAs, compared to non-PDA control cells (HPDE and 293T). Data are shown as mean ± SD. Comparison of groups carried out by and one-way ANOVA (**A**). ****p<0.0001, ***p<0.001, **p<0.01, *p<0.05.

**Fig. S3.**
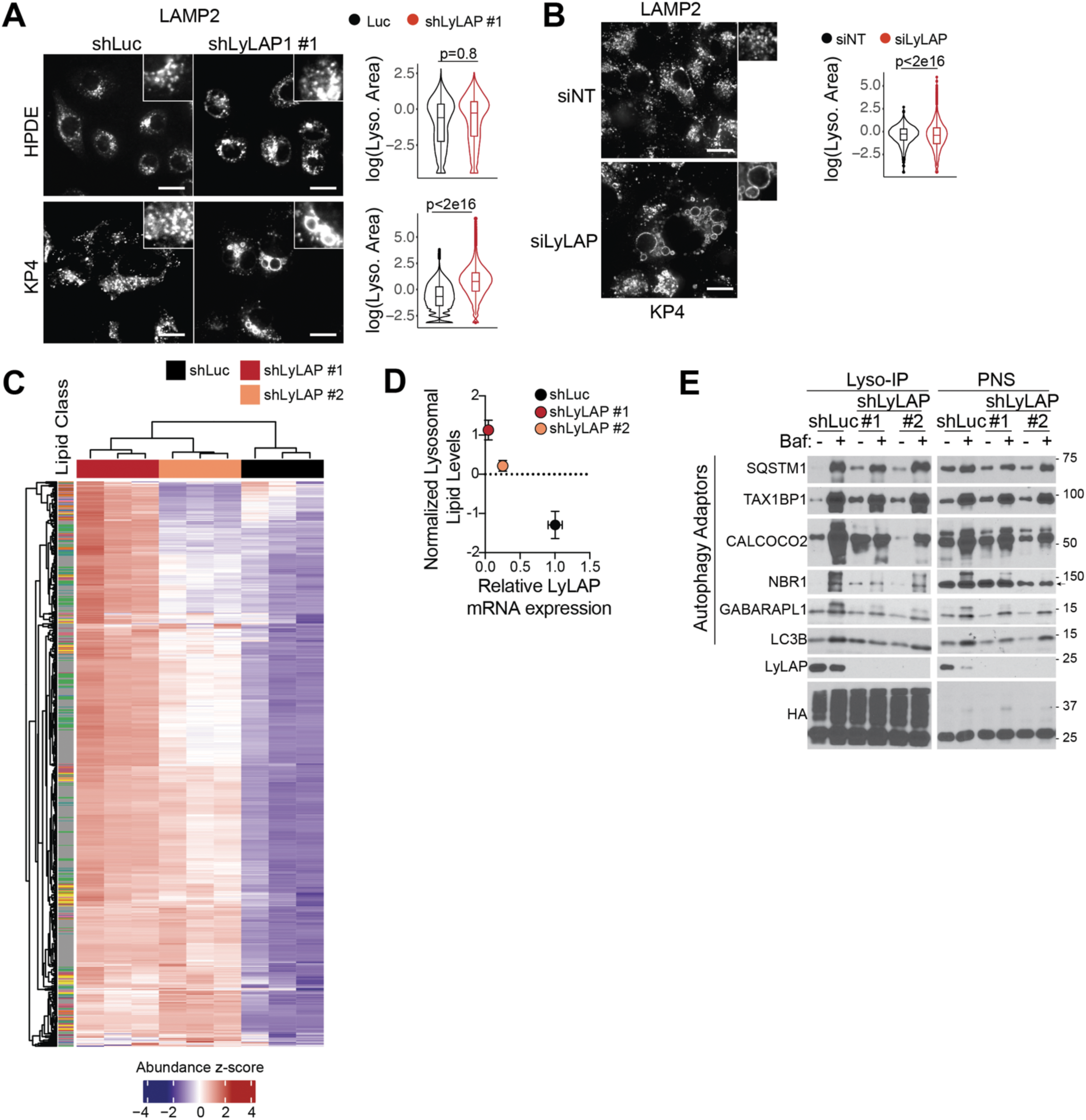
LyLAP depletion causes lysosomal dysfunction in PDA. **A.** (*left*) Representative immunofluorescence images of LAMP2-stained HPDE (non-PDA) and KP4 (PDA) cells transduced with the indicated shRNAs. (*right*) Quantification of lysosome area for the respective cell lines and shRNA conditions on the left. Scale bars, 10 μm. **B.** (*left*) Representative immunofluorescence images of LAMP2-stained KP4 cells transfected with non-targeting siRNA pool (siNT), or with LyLAP-targeting siRNA pool. (*right*) Quantification of lysosome area for the respective cell lines and siRNA conditions on the left. Scale bars, 10 μm. **C.** (related to the untargeted lipidomics experiment described in Fig. 2H): Heatmap displaying normalized abundance of lipid species quantified from KP4 lysosome immunoprecipitates, following cell treatment with the indicated shRNAs. **D.** Normalized lysosomal lipid concentrations in KP4 cells treated with the indicated shRNAs, plotted against LyLAP mRNA expression measured by qPCR. Normalized lipid concentrations generated from respective lipid abundance values shown in **C**. **E.** Immunoblot of autophagy adaptors in lysosomal lysates immunopurified from KP4 cells, following treatment with the indicated shRNAs and with either DMSO (vehicle) or BafA1. Data are shown as mean ± SD. Comparison of groups carried out by two-tailed, unpaired t-test (**A, B**).

**Fig. S4.**
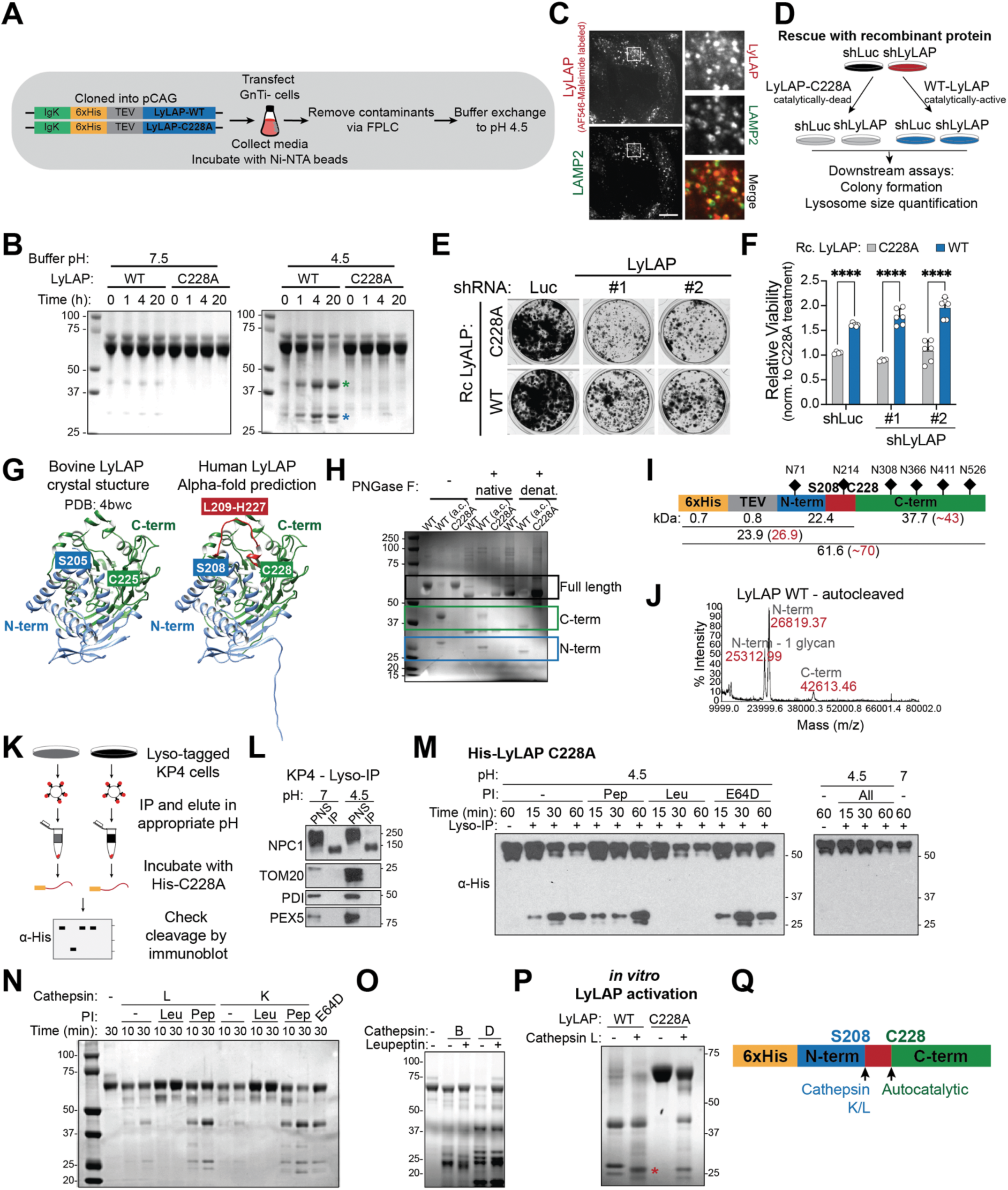
LyLAP is proteolytically activated within lysosomes. **A.** Schematic of recombinant LyLAP purification strategy in 293 GnTI-cells using IgK secretion signal-tagged protein for subsequent purification from media supernatant. **B.** Coomassie staining showing time-dependent autocatalytic cleavage of wild-type (WT), but not catalytic-dead C228A mutant, LyLAP at pH 4.5 (*right*), but not at neutral pH (*left*). Blue and green asterisks indicate N-and C-termini of the autocatalytically cleaved protein, respectively. **C.** Representative immunofluorescence image of KP4 cells treated with AF546-maleimide-labeled recombinant WT LyLAP and stained with an antibody for the lysosomal marker, LAMP2. Scale bar, 10 μm. **D.** Schematic depicting experimental set-up for rescue of LyLAP-KD phenotypes with recombinant protein. KP4 cells were treated with either shLuc or shLyLAP and supplemented with either WT LyLAP or catalytically dead C228A mutant, prior to seeding for subsequent assays –colony formation and lysosomal size quantification. **E.** Representative images of colony formation assay of KP4 cells treated with indicated shRNAs and supplemented with WT or C228A LyLAP. **F.** Crystal violet dye absorbance from colony formation assay in (**E**) to assess relative viability. For each respective shRNA condition, data were normalized to the C228A-treated samples. Data are shown as mean ± SD. Comparison of groups carried out by two-way ANOVA. ****p<0.0001. **G.** Comparison of the bovine LyLAP crystal structure (PDB: 4bwc) with the AlphaFold prediction for human LyLAP. Blue and green domains refer to the N-and C-terminus of the autocatalytically cleaved protein, respectively; red loop (L209-H227) on human LyLAP prediction refers to the loop missing from the bovine crystal structure. **H.** Coomassie staining of PNGase F-deglycosylated full-length LyLAP, autocatalytically cleaved (WT a.c.) LyLAP and C228A mutant by treatment with in native or denaturing conditions. **I.** Schematic of recombinant LyLAP with expected molecular weights and mapping of N-glycosylation sites. Molecular weights in red denote values with expected glycans added. **J.** MALDI-TOF mass spectra of recombinant WT LyLAP autocatalytically cleaved via overnight incubation at pH 4.5. **K.** Schematic depicting experimental set-up for incubation of recombinant, His-tagged LyLAP C228A mutant with lysosomal lysates immunopurified from KP4 cells for the detection of a secondary cleavage site on the protein. **L.** Immunoblot verification of lysosome immunoprecipitates from KP4 cells, following elution at pH 7 versus pH 4.5 (lysates used in **M**). **M.** Immunoblot analysis for the time-dependent cleavage of recombinant, His-tagged LyLAP C228A mutant upon incubation with KP4-derived lysosomal lysates in the presence of the indicated protease inhibitors, Leupeptin (Leu), Pepstatin (Pep), E64D or neutral pH. **N.** Coomassie of *in vitro* cleaved recombinant LyLAP C228A mutant when incubated with Cathepsin L or Cathepsin K recombinant proteins. Indicated protease inhibitors used as controls. **O.** Coomassie staining of *in vitro*-cleaved recombinant LyLAP C228A mutant when incubated with Cathepsin B or D recombinant proteins. Leupeptin was used as control. **P.** Coomassie staining of WT and C228A LyLAP proteolytically activated by sequential autocatalytic and recombinant Cathepsin L-mediated cleavage *in vitro*. Red asterisk indicates further trimming of the WT N-terminal fragment by Cathepsin L. **Q.** Model of LyLAP activation by autocatalytic cleavage at C228 followed by cathepsin L or K-mediated cleavage at S208.

**Fig. S5.**
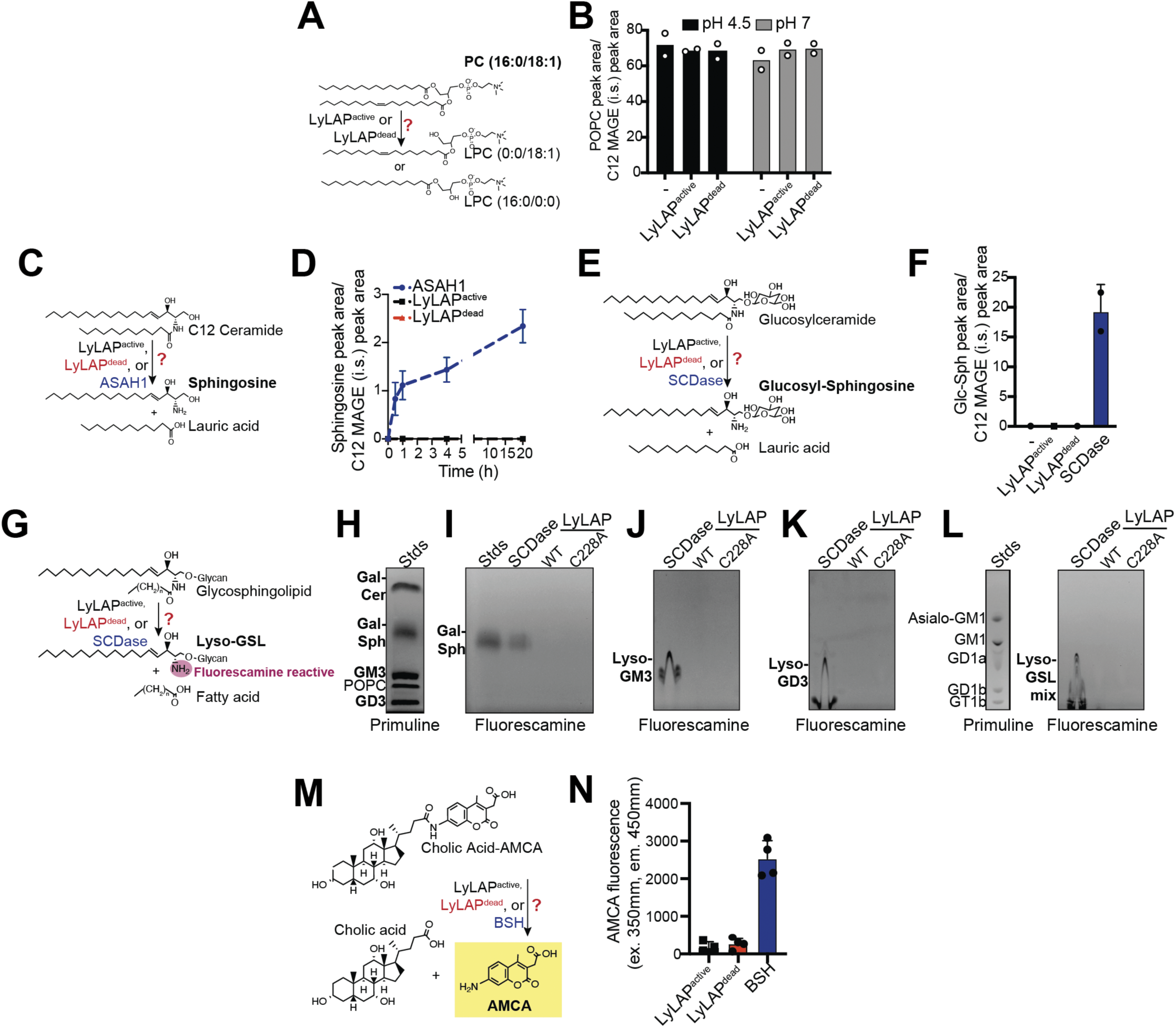
LyLAP is not a lipid hydrolase. **A.** Schematic of assay for putative phospholipase B activity of LyLAP^active^ with 16:0/18:1 PC substrate and 18:1 LPC or 16:0 LPC products. LyLAP^dead^ used as negative control for this assay. 16:0/18:1 PC (in bold) was detected by LC-MS. **B.** 16:0/18:1 PC peak area, normalized to C12-mono-acyl-glycerin-ether (C12 MAGE) internal standard per run, upon 24 h incubation with either LyLAP^active^ or LyLAP^dead^ at either pH 4.5 or pH 7. **C.** Schematic of assay for putative acid ceramidase activity of LyLAP^active^ with C12-ceramide substrate and sphingosine or lauric acid products. LyLAP^dead^ and ASAH1 used as negative and positive controls, respectively, for this activity. Sphingosine (in bold) detected by LC-MS. **D.** Sphingosine peak area over time (normalized to C12 MAGE internal standard per run) over 20 h incubation period at pH 4.5 with either ASAH1, LyLAP^active^ or LyLAP^dead^. **E.** Schematic of assay for putative glycosphingolipid deacylase activity of LyLAP^active^ with C12-glucosylceramide substrate and glucosylsphingosine and lauric acid products. LyLAP^dead^ and bacterial sphingolipid ceramide N-deacylase (SCDase) used as negative and positive controls, respectively, for this activity. Glucosylsphingosine (in bold) detected by LC-MS. **F.** Glucosylsphingosine peak area (normalized to C12 MAGE internal standard per run) upon 24 h incubation with either LyLAP^active^, LyLAP^dead^ or SCDase. **G.** Schematic of assay for putative complex glycosphingolipid deacylase activity of LyLAP^active^ with various glycosphingolipid substrates and lyso-glycosphingolipid (lyso-GSL) and the appropriate fatty acid products. Primary amine of Lyso-GSLs (in bold) react with fluorescamine and was visualized by thin-layer chromatography (TLC). LyLAP^dead^ and bacterial SCDase used as negative and positive controls, respectively. **H.** TLC of substrate glycolipids, galactosyl-ceramide (Gal-Cer), galactosyl-sphingosine (Gal-Sph), GM3 and GD3, as detected by primuline staining. **I-K.** TLC of galactosyl-sphingosine (Gal-Sph) (**I**), Lyso-GM3 (**J**), and lyso-GD3 (**K**) product formation upon incubation of Gal-Cer, GM3, or GD3, respectively, with bacterial SCDase (positive control), LyLAP^active^ or LyLAP^dead^ as detected by fluorescamine staining. **L.** TLC of ganglioside standard mix detected by primuline staining (left) and product formation upon incubation with either SCDase, LyLAP^active^ or LyLAP^dead^ by fluorescamine staining (right). **M.** Schematic of assay for putative bile salt hydrolase activity of LyLAP^active^ with cholate-AMCA substrate and cholic acid and AMCA as products. LyLAP^dead^ and bacterial bile salt hydrolase (BSH) used as negative and positive controls, respectively, for this activity. AMCA is fluorescent upon release and detected by plate reader. **N.** Maximum AMCA fluorescence upon incubation with either BSH, LyLAP^active^ or LyLAP^dead^ over 20 min incubation period at pH 4.5.

**Fig. S6.**
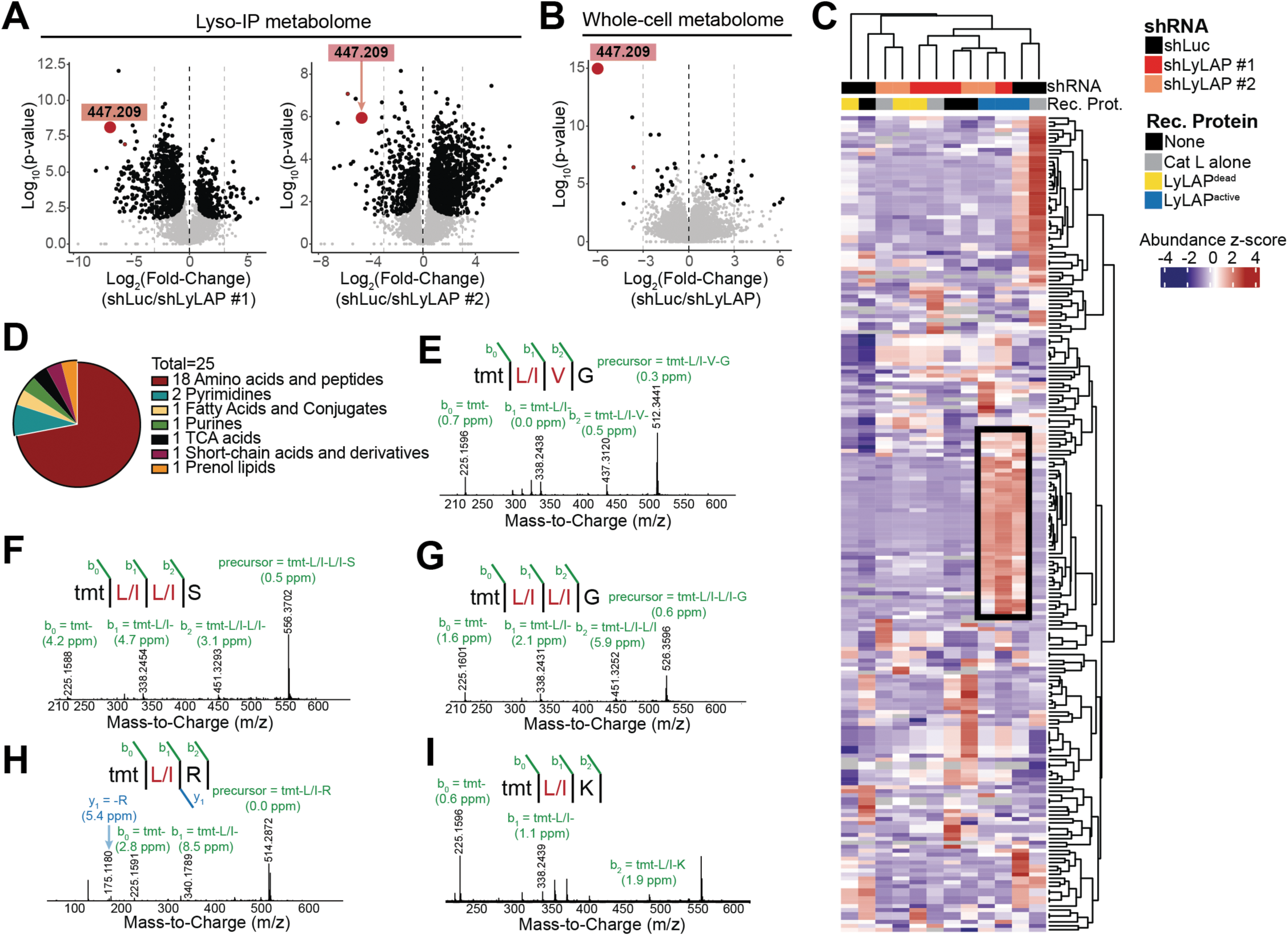
LyLAP substrates are peptides. **A.** Volcano plots from untargeted metabolomic analysis of lysosomes immunoprecipitated from Lyso-tagged KP4 cells harboring either shLyLAP #1 (left) or shLyLAP #2 (right), each compared to shLuc control. The highlighted m/z ratio is a putative substrate. **B.** Volcano plot from untargeted metabolomic analysis of whole cell extracts from KP4 cells treated with shLyLAP versus shLuc control. Highlighted m/z ratio is a putative substrate. **C.** Heatmap of verified metabolites from all the untargeted metabolomic samples (as described in Fig. 3B) identifies a cluster of small metabolites (black box) that are only enriched when lysates are treated with recombinant LyLAP^active^. **D.** Composition analysis of boxed (enriched upon LyLAP^active^ treatment) cluster of metabolites from (**C**). **E-I.** MS2 spectra of TMT-derivatized peptides that follow pattern of accumulation upon LyLAP-KD and depletion upon LyLAP^active^ treatment (from Fig. 4E).

**Fig. S7.**
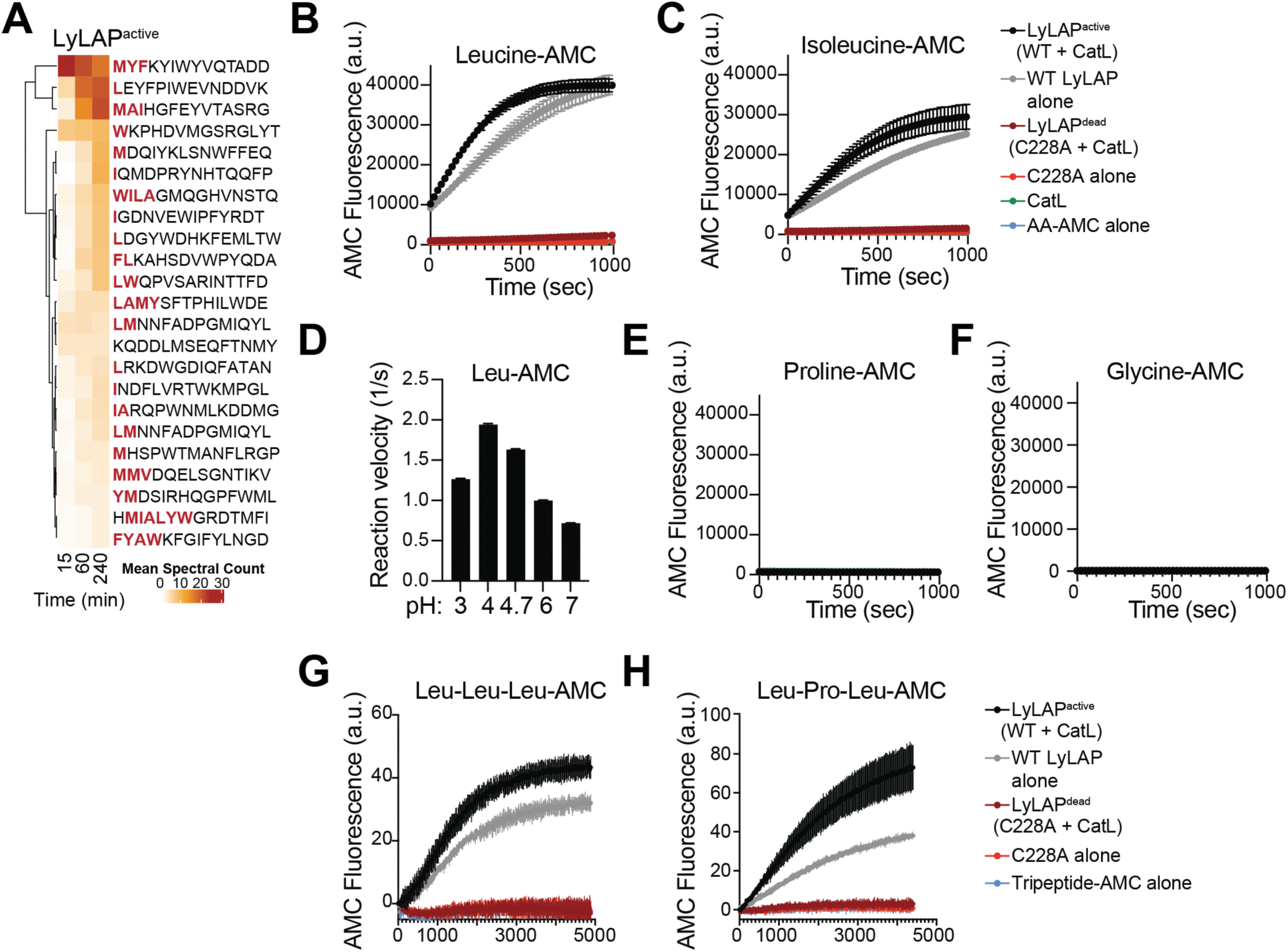
LyLAP is a processive monoaminopeptidase for hydrophobic amino acids. **A.** Heatmap of mean spectral counts indicating peptide cleavage events from MSP-MS library for LyLAP^active^ over incubation time. Hydrophobic N-terminal residues on each peptide highlighted in red. **B, C.** AMC fluorescence over time upon incubation of Leu-AMC (**B**), Ile-AMC **(C)**, with LyLAP^active^, autocatalytically cleaved LyLAP (WT LyLAP alone), LyLAP^dead^, C228A mutant alone, cathepsin L alone (CatL) or no protein (AA-AMC alone). **D.** Reaction rate curve of LyLAP^active^ against Leu-AMC substrate over pH range from 3 to 7. **E, F.** AMC fluorescence over time upon incubation of Pro-AMC (**E**), Gly-AMC **(F)**, with LyLAP^active^, autocatalytically cleaved LyLAP (WT LyLAP alone), LyLAP^dead^, C228A mutant alone, cathepsin L alone (CatL) or no protein (AA-AMC alone). **G, H.** AMC fluorescence over time upon incubation of Leu-Leu-Leu-AMC (**G**), Leu-Pro-Leu-AMC **(H)**, with LyLAP^active^, autocatalytically cleaved LyLAP (WT LyLAP alone), LyLAP^dead^, C228A mutant alone or no protein (Tripeptide-AMC alone).

**Fig. S8.**
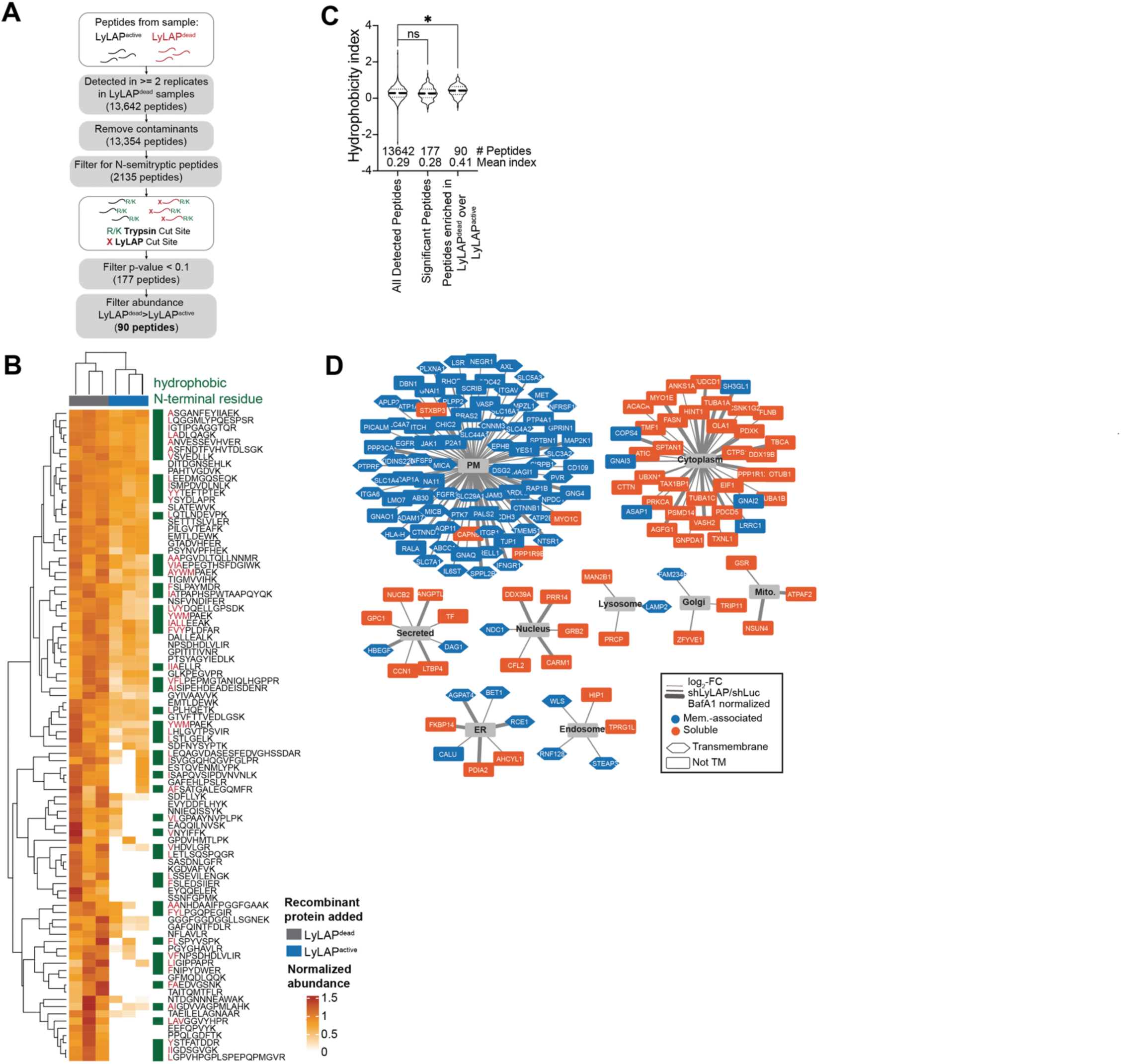
LyLAP enables transmembrane protein degradation in PDA lysosomes. **A.** Schematic of analytical pipeline for N-semitryptic peptide determination (related to experiment described in Fig. 4A). **B.** Heatmap of normalized abundance values of LyLAP substrate peptides from experiment described in Fig. 4A, using the analytical method shown in (**A**). **C.** Hydrophobicity index (by Abraham-Leo scale) of LyLAP substrate peptides (enriched in LyLAP^dead^ over LyLAP^active^) compared to all detected peptides. Comparison of groups carried out by one-way ANOVA. *p<0.05, ns=not significant. **D.** Network analysis of LyLAP substrate proteins as determined from lysosomal proteomics analysis described in Fig. 4G. Membrane-associated and soluble proteins annotated in blue and orange, respectively; transmembrane versus non-transmembrane protein annotated by hexagonal versus rectangular node shape, respectively. PM, plasma membrane; ER, endoplasmic reticulum, Mito., mitochondria.

**Fig. S9.**
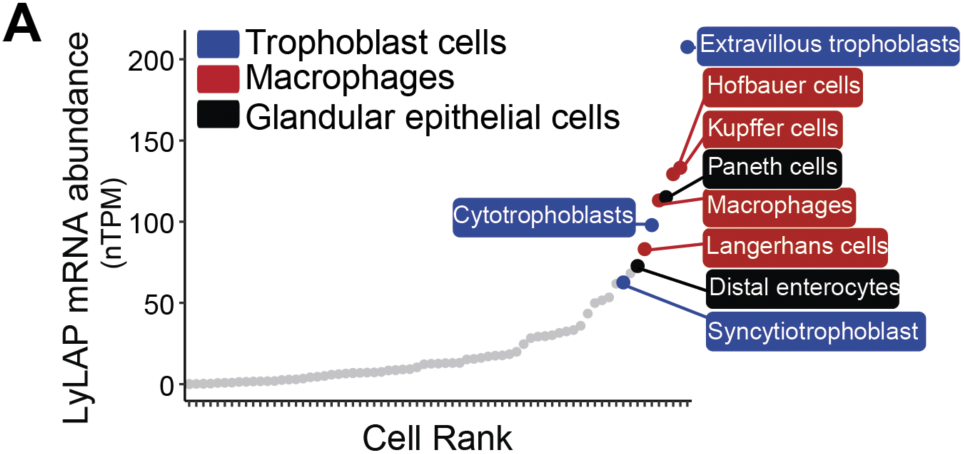
LyLAP expression across non-transformed cell types. **A.** Waterfall plot of PLBD1 mRNA expression from single-cell RNAseq data (from Human Protein Atlas).

**Data S1. (separate file)**

Proteomic analysis of lysosomes immunoprecipitated from 8988T versus HPDE cells.

**Data S2. (separate file)**

Untargeted lipidomic analysis of lysosomes immunoprecipitated from KP4 cells following treatment with shLuc, shLyLAP #1 or shLyLAP #2.

**Data S3. (separate file)**

Proteomic analysis of lysosomes immunoprecipitated from KP4 cells following treatment with shLuc, shLyLAP #1 or shLyLAP #2 and further treated with either Bafilomycin A1 or DMSO vehicle control.

**Data S4. (separate file)**

Untargeted metabolomic analysis of lysosomes immunoprecipitated from KP4 cells following treatment with shLuc, shLyLAP #1 or shLyLAP #2.

**Data S5. (separate file)**

Untargeted metabolomic analysis of lysosomes immunoprecipitated from KP4 cells following treatment with shLuc, shLyLAP #1 or shLyLAP #2 followed by add-back of either recombinant LyLAP^active^, LyLAP^dead^ or cathepsin L (control). Targeted subset of metabolites only.

**Data S6. (separate file)**

Peptide abundances from MS2 analysis from lysosomes immunoprecipitated from KP4 cells following treatment with shLuc, shLyLAP #1 or shLyLAP #2 followed by add-back of either recombinant LyLAP^active^, LyLAP^dead^ or cathepsin L (control). Targeted subset of metabolites only.

**Data S7. (separate file)**

MSP-MS analysis of recombinant LyLAP^active^.

**Data S8. (separate file)**

Semi-tryptic peptidomic analysis of lysosomes immunoprecipitated from KP4 cells following treatment with shLyLAP #1 followed by add-back of either recombinant LyLAP^active^ or LyLAP^dead^. Hydrophobicity indices of each peptide determined by the Abraham-Leo scale.

**Data S9. (separate file)**

Peptidomic analysis of lysosomes immunoprecipitated from KP4 cells following treatment with shLuc or shLyLAP #1. Hydrophobicity indices of each peptide determined by the Abraham-Leo scale.

## References and Notes

1. L. Dobson, I. Reményi, G. E. Tusnády, The human transmembrane proteome. Biol Direct 10, 31 (2015).

2. L. Fagerberg, K. Jonasson, G. von Heijne, M. Uhlén, L. Berglund, Prediction of the human membrane proteome. Proteomics 10, 1141–1149 (2010).

3. M. A. Jambrich, G. E. Tusnady, L. Dobson, How AlphaFold2 shaped the structural coverage of the human transmembrane proteome. Sci Rep 13, 20283 (2023).

4. J. Pei, Q. Cong, AFTM: a database of transmembrane regions in the human proteome predicted by AlphaFold. Database (Oxford) 2023, baad008 (2023).

5. J. A. MacGurn, P.-C. Hsu, S. D. Emr, Ubiquitin and membrane protein turnover: from cradle to grave. Annu Rev Biochem 81, 231–259 (2012).

6. C. Raiborg, H. Stenmark, The ESCRT machinery in endosomal sorting of ubiquitylated membrane proteins. Nature 458, 445–452 (2009).

7. H. Schulze, T. Kolter, K. Sandhoff, Principles of lysosomal membrane degradation: Cellular topology and biochemistry of lysosomal lipid degradation. Biochim Biophys Acta 1793, 674–683 (2009).

8. C. Settembre, R. M. Perera, Lysosomes as coordinators of cellular catabolism, metabolic signalling and organ physiology. Nat Rev Mol Cell Biol 25, 223–245 (2024).

9. N. De Franceschi, H. Hamidi, J. Alanko, P. Sahgal, J. Ivaska, Integrin traffic -the update. J Cell Sci 128, 839–852 (2015).

10. H. E. Grecco, M. Schmick, P. I. H. Bastiaens, Signaling from the living plasma membrane. Cell 144, 897–909 (2011).

11. A. Sorkin, M. von Zastrow, Endocytosis and signalling: intertwining molecular networks. Nat Rev Mol Cell Biol 10, 609–622 (2009).

12. S. Sigismund, L. Lanzetti, G. Scita, P. P. Di Fiore, Endocytosis in the context-dependent regulation of individual and collective cell properties. Nat Rev Mol Cell Biol 22, 625–643 (2021).

13. C. H. Lin, J. A. MacGurn, T. Chu, C. J. Stefan, S. D. Emr, Arrestin-related ubiquitin-ligase adaptors regulate endocytosis and protein turnover at the cell surface. Cell 135, 714–725 (2008).

14. V. Turk, V. Stoka, O. Vasiljeva, M. Renko, T. Sun, B. Turk, D. Turk, Cysteine cathepsins: from structure, function and regulation to new frontiers. Biochim Biophys Acta 1824, 68–88 (2012).

15. C. Commisso, S. M. Davidson, R. G. Soydaner-Azeloglu, S. J. Parker, J. J. Kamphorst, S. Hackett, E. Grabocka, M. Nofal, J. A. Drebin, C. B. Thompson, J. D. Rabinowitz, C. M. Metallo, M. G. Vander Heiden, D. Bar-Sagi, Macropinocytosis of protein is an amino acid supply route in Ras-transformed cells. Nature 497, 633–637 (2013).

16. R. M. Perera, S. Stoykova, B. N. Nicolay, K. N. Ross, J. Fitamant, M. Boukhali, J. Lengrand, V. Deshpande, M. K. Selig, C. R. Ferrone, J. Settleman, G. Stephanopoulos, N. J. Dyson, R. Zoncu, S. Ramaswamy, W. Haas, N. Bardeesy, Transcriptional control of autophagy-lysosome function drives pancreatic cancer metabolism. Nature 524, 361–365 (2015).

17. S. Yang, X. Wang, G. Contino, M. Liesa, E. Sahin, H. Ying, A. Bause, Y. Li, J. M. Stommel, G. Dell’antonio, J. Mautner, G. Tonon, M. Haigis, O. S. Shirihai, C. Doglioni, N. Bardeesy, A. C. Kimmelman, Pancreatic cancers require autophagy for tumor growth. Genes Dev 25, 717–729 (2011).

18. W. Palm, Y. Park, K. Wright, N. N. Pavlova, D. A. Tuveson, C. B. Thompson, The Utilization of Extracellular Proteins as Nutrients Is Suppressed by mTORC1. Cell 162, 259–270 (2015).

19. J. B. Casaletto, A. I. McClatchey, Spatial regulation of receptor tyrosine kinases in development and cancer. Nat Rev Cancer 12, 387–400 (2012).

20. S. L. Sønder, S. C. Häger, A. S. B. Heitmann, L. B. Frankel, C. Dias, A. C. Simonsen, J. Nylandsted, Restructuring of the plasma membrane upon damage by LC3-associated macropinocytosis. Sci Adv 7, eabg1969 (2021).

21. K. Rose, T. Jepson, S. Shukla, A. Maya-Romero, M. Kampmann, K. Xu, J. H. Hurley, Tau fibrils induce nanoscale membrane damage and nucleate cytosolic tau at lysosomes. Proc Natl Acad Sci U S A 121, e2315690121 (2024).

22. J. J. Chen, D. L. Nathaniel, P. Raghavan, M. Nelson, R. Tian, E. Tse, J. Y. Hong, S. K. See, S.-A. Mok, M. Y. Hein, D. R. Southworth, L. T. Grinberg, J. E. Gestwicki, M. D. Leonetti, M. Kampmann, Compromised function of the ESCRT pathway promotes endolysosomal escape of tau seeds and propagation of tau aggregation. J Biol Chem 294, 18952–18966 (2019).

23. C. Bussi, A. Mangiarotti, C. Vanhille-Campos, B. Aylan, E. Pellegrino, N. Athanasiadi, A. Fearns, A. Rodgers, T. M. Franzmann, A. Šarić, R. Dimova, M. G. Gutierrez, Stress granules plug and stabilize damaged endolysosomal membranes. Nature 623, 1062–1069 (2023).

24. I. Maejima, A. Takahashi, H. Omori, T. Kimura, Y. Takabatake, T. Saitoh, A. Yamamoto, M. Hamasaki, T. Noda, Y. Isaka, T. Yoshimori, Autophagy sequesters damaged lysosomes to control lysosomal biogenesis and kidney injury. EMBO J 32, 2336–2347 (2013).

25. M. Radulovic, K. O. Schink, E. M. Wenzel, V. Nähse, A. Bongiovanni, F. Lafont, H. Stenmark, ESCRT-mediated lysosome repair precedes lysophagy and promotes cell survival. EMBO J 37, e99753 (2018).

26. M. L. Skowyra, P. H. Schlesinger, T. V. Naismith, P. I. Hanson, Triggered recruitment of ESCRT machinery promotes endolysosomal repair. Science 360, eaar5078 (2018).

27. J. X. Tan, T. Finkel, A phosphoinositide signalling pathway mediates rapid lysosomal repair. Nature 609, 815–821 (2022).

28. H. Yang, J. X. Tan, Lysosomal quality control: molecular mechanisms and therapeutic implications. Trends Cell Biol 33, 749–764 (2023).

29. R. Zoncu, R. M. Perera, Built to last: lysosome remodeling and repair in health and disease. Trends Cell Biol 32, 597–610 (2022).

30. S. H. White, W. C. Wimley, Hydrophobic interactions of peptides with membrane interfaces. Biochim Biophys Acta 1376, 339–352 (1998).

31. S. M. Banik, K. Pedram, S. Wisnovsky, G. Ahn, N. M. Riley, C. R. Bertozzi, Lysosome-targeting chimaeras for degradation of extracellular proteins. Nature 584, 291–297 (2020).

32. G. Ahn, N. M. Riley, R. A. Kamber, S. Wisnovsky, S. Moncayo von Hase, M. C. Bassik, S. M. Banik, C. R. Bertozzi, Elucidating the cellular determinants of targeted membrane protein degradation by lysosome-targeting chimeras. Science 382, eadf6249 (2023).

33. M. S. Haney, C. J. Bohlen, D. W. Morgens, J. A. Ousey, A. A. Barkal, C. K. Tsui, B. K. Ego, R. Levin, R. A. Kamber, H. Collins, A. Tucker, A. Li, D. Vorselen, L. Labitigan, E. Crane, E. Boyle, L. Jiang, J. Chan, E. Rincón, W. J. Greenleaf, B. Li, M. P. Snyder, I. L. Weissman, J. A. Theriot, S. R. Collins, B. A. Barres, M. C. Bassik, Identification of phagocytosis regulators using magnetic genome-wide CRISPR screens. Nat Genet 50, 1716–1727 (2018).

34. B. Lindner, E. Martin, M. Steininger, A. Bundalo, M. Lenter, J. Zuber, M. Schuler, A genome-wide CRISPR/Cas9 screen to identify phagocytosis modulators in monocytic THP-1 cells. Sci Rep 11, 12973 (2021).

35. T. S. P. Heng, M. W. Painter, Immunological Genome Project Consortium, The Immunological Genome Project: networks of gene expression in immune cells. Nat Immunol 9, 1091–1094 (2008).

36. M. Abu-Remaileh, G. A. Wyant, C. Kim, N. N. Laqtom, M. Abbasi, S. H. Chan, E. Freinkman, D. M. Sabatini, Lysosomal metabolomics reveals V-ATPase-and mTOR-dependent regulation of amino acid efflux from lysosomes. Science 358, 807–813 (2017).

37. O. B. Davis, H. R. Shin, C.-Y. Lim, E. Y. Wu, M. Kukurugya, C. F. Maher, R. M. Perera, M. P. Ordonez, R. Zoncu, NPC1-mTORC1 Signaling Couples Cholesterol Sensing to Organelle Homeostasis and Is a Targetable Pathway in Niemann-Pick Type C. Dev Cell 56, 260–276.e7 (2021).

38. S. Gupta, J. Yano, V. Mercier, H. H. Htwe, H. R. Shin, G. Rademaker, Z. Cakir, T. Ituarte, K. W. Wen, G. E. Kim, R. Zoncu, A. Roux, D. W. Dawson, R. M. Perera, Lysosomal retargeting of Myoferlin mitigates membrane stress to enable pancreatic cancer growth. Nat Cell Biol 23, 232–242 (2021).

39. R. K. Amaravadi, A. C. Kimmelman, J. Debnath, Targeting Autophagy in Cancer: Recent Advances and Future Directions. Cancer Discov 9, 1167–1181 (2019).

40. K. L. Bryant, C. A. Stalnecker, D. Zeitouni, J. E. Klomp, S. Peng, A. P. Tikunov, V. Gunda, M. Pierobon, A. M. Waters, S. D. George, G. Tomar, B. Papke, G. A. Hobbs, L. Yan, T. K. Hayes, J. N. Diehl, G. D. Goode, N. V. Chaika, Y. Wang, G.-F. Zhang, A. K. Witkiewicz, E. S. Knudsen, E. F. Petricoin, P. K. Singh, J. M. Macdonald, N. L. Tran, C. A. Lyssiotis, H. Ying, A. C. Kimmelman, A. D. Cox, C. J. Der, Combination of ERK and autophagy inhibition as a treatment approach for pancreatic cancer. Nat Med 25, 628–640 (2019).

41. C. G. Kinsey, S. A. Camolotto, A. M. Boespflug, K. P. Guillen, M. Foth, A. Truong, S. S. Schuman, J. E. Shea, M. T. Seipp, J. T. Yap, L. D. Burrell, D. H. Lum, J. R. Whisenant, G. W. Gilcrease, C. C. Cavalieri, K. M. Rehbein, S. L. Cutler, K. E. Affolter, A. L. Welm, B. E. Welm, C. L. Scaife, E. L. Snyder, M. McMahon, Protective autophagy elicited by RAF→MEK→ERK inhibition suggests a treatment strategy for RAS-driven cancers. Nat Med 25, 620–627 (2019).

42. R. Zoncu, R. M. Perera, Emerging roles of the MiT/TFE factors in cancer. Trends Cancer 9, 817–827 (2023).

43. H. Repo, E. Kuokkanen, E. Oksanen, A. Goldman, P. Heikinheimo, Is the bovine lysosomal phospholipase B-like protein an amidase? Proteins 82, 300–311 (2014).

44. S. Xu, L. Zhao, A. Larsson, P. Venge, The identification of a phospholipase B precursor in human neutrophils. FEBS J 276, 175–186 (2009).

45. H. Wang, J. L. Rubinstein, CryoEM of V-ATPases: Assembly, disassembly, and inhibition. Curr Opin Struct Biol 80, 102592 (2023).

46. A. H. Ponsford, T. A. Ryan, A. Raimondi, E. Cocucci, S. A. Wycislo, F. Fröhlich, L. E. Swan, M. Stagi, Live imaging of intra-lysosome pH in cell lines and primary neuronal culture using a novel genetically encoded biosensor. Autophagy 17, 1500–1518 (2021).

47. V. Rogov, V. Dötsch, T. Johansen, V. Kirkin, Interactions between autophagy receptors and ubiquitin-like proteins form the molecular basis for selective autophagy. Mol Cell 53, 167– 178 (2014).

48. G. Zaffagnini, S. Martens, Mechanisms of Selective Autophagy. J Mol Biol 428, 1714– 1724 (2016).

49. S. Zellner, M. Schifferer, C. Behrends, Systematically defining selective autophagy receptor-specific cargo using autophagosome content profiling. Mol Cell 81, 1337–1354.e8 (2021).

50. C. Oinonen, J. Rouvinen, Structural comparison of Ntn-hydrolases. Protein Sci 9, 2329– 2337 (2000).

51. K. Tunyasuvunakool, J. Adler, Z. Wu, T. Green, M. Zielinski, A. Žídek, A. Bridgland, A. Cowie, C. Meyer, A. Laydon, S. Velankar, G. J. Kleywegt, A. Bateman, R. Evans, A. Pritzel, M. Figurnov, O. Ronneberger, R. Bates, S. A. A. Kohl, A. Potapenko, A. J. Ballard, B. Romera-Paredes, S. Nikolov, R. Jain, E. Clancy, D. Reiman, S. Petersen, A. W. Senior, K. Kavukcuoglu, E. Birney, P. Kohli, J. Jumper, D. Hassabis, Highly accurate protein structure prediction for the human proteome. Nature 596, 590–596 (2021).

52. B. Breiden, K. Sandhoff, Lysosomal Glycosphingolipid Storage Diseases. Annu Rev Biochem 88, 461–485 (2019).

53. I. P. Heremans, F. Caligiore, I. Gerin, M. Bury, M. Lutz, J. Graff, V. Stroobant, D. Vertommen, A. A. Teleman, E. Van Schaftingen, G. T. Bommer, Parkinson’s disease protein PARK7 prevents metabolite and protein damage caused by a glycolytic metabolite. Proc Natl Acad Sci U S A 119, e2111338119 (2022).

54. P. J. Rohweder, Z. Jiang, B. M. Hurysz, A. J. O’Donoghue, C. S. Craik, Multiplex substrate profiling by mass spectrometry for proteases. Methods Enzymol 682, 375–411 (2023).

55. D. Eisenberg, Three-dimensional structure of membrane and surface proteins. Annu Rev Biochem 53, 595–623 (1984).

56. D. Eisenberg, R. M. Weiss, T. C. Terwilliger, The helical hydrophobic moment: a measure of the amphiphilicity of a helix. Nature 299, 371–374 (1982).

57. V. H. Lobert, A. Brech, N. M. Pedersen, J. Wesche, A. Oppelt, L. Malerød, H. Stenmark, Ubiquitination of alpha 5 beta 1 integrin controls fibroblast migration through lysosomal degradation of fibronectin-integrin complexes. Dev Cell 19, 148–159 (2010).

58. H. Heerklotz, Interactions of surfactants with lipid membranes. Q Rev Biophys 41, 205–264 (2008).

59. U. Repnik, M. Borg Distefano, M. T. Speth, M. Y. W. Ng, C. Progida, B. Hoflack, J. Gruenberg, G. Griffiths, L-leucyl-L-leucine methyl ester does not release cysteine cathepsins to the cytosol but inactivates them in transiently permeabilized lysosomes. J Cell Sci 130, 3124–3140 (2017).

60. M. B. Ulmschneider, J. C. Smith, J. P. Ulmschneider, Peptide partitioning properties from direct insertion studies. Biophys J 98, L60–62 (2010).

61. P. Gahlot, B. Kravic, G. Rota, J. van den Boom, S. Levantovsky, N. Schulze, E. Maspero, S. Polo, C. Behrends, H. Meyer, Lysosomal damage sensing and lysophagy initiation by SPG20-ITCH. Mol Cell 84, 1556–1569.e10 (2024).

62. D. L. Thiele, P. E. Lipsky, Mechanism of L-leucyl-L-leucine methyl ester-mediated killing of cytotoxic lymphocytes: dependence on a lysosomal thiol protease, dipeptidyl peptidase I, that is enriched in these cells. Proc Natl Acad Sci U S A 87, 83–87 (1990).

63. R. M. Perera, N. Bardeesy, Pancreatic Cancer Metabolism: Breaking It Down to Build It Back Up. Cancer Discov 5, 1247–1261 (2015).

64. M. Matsui, J. H. Fowler, L. L. Walling, Leucine aminopeptidases: diversity in structure and function. Biol Chem 387, 1535–1544 (2006).

65. H. Meyer, B. Kravic, The Endo-Lysosomal Damage Response. Annu Rev Biochem, doi: 10.1146/annurev-biochem-030222-102505 (2024).

66. C. Pechincha, S. Groessl, R. Kalis, M. de Almeida, A. Zanotti, M. Wittmann, M. Schneider, R. P. de Campos, S. Rieser, M. Brandstetter, A. Schleiffer, K. Müller-Decker, D. Helm, S. Jabs, D. Haselbach, M. K. Lemberg, J. Zuber, W. Palm, Lysosomal enzyme trafficking factor LYSET enables nutritional usage of extracellular proteins. Science 378, eabn5637 (2022).

67. K. Lakomek, A. Dickmanns, M. Kettwig, H. Urlaub, R. Ficner, T. Lübke, Initial insight into the function of the lysosomal 66.3 kDa protein from mouse by means of X-ray crystallography. BMC Struct Biol 9, 56 (2009).

68. T. Takahashi, A. H. Dehdarani, J. Tang, Porcine spleen cathepsin H hydrolyzes oligopeptides solely by aminopeptidase activity. J Biol Chem 263, 10952–10957 (1988).

69. J. Behnke, J. Schneppenheim, F. Koch-Nolte, F. Haag, P. Saftig, B. Schröder, Signal-peptide-peptidase-like 2a (SPPL2a) is targeted to lysosomes/late endosomes by a tyrosine motif in its C-terminal tail. FEBS Lett 585, 2951–2957 (2011).

70. A. Manuyakorn, R. Paulus, J. Farrell, N. A. Dawson, S. Tze, G. Cheung-Lau, O. J. Hines, H. Reber, D. B. Seligson, S. Horvath, S. K. Kurdistani, C. Guha, D. W. Dawson, Cellular histone modification patterns predict prognosis and treatment response in resectable pancreatic adenocarcinoma: results from RTOG 9704. J Clin Oncol 28, 1358–1365 (2010).

71. D. R. Stirling, M. J. Swain-Bowden, A. M. Lucas, A. E. Carpenter, B. A. Cimini, A. Goodman, CellProfiler 4: improvements in speed, utility and usability. BMC Bioinformatics 22, 433 (2021).

72. K. S. Beckwith, M. S. Beckwith, S. Ullmann, R. S. Sætra, H. Kim, A. Marstad, S. E. Åsberg, T. A. Strand, M. Haug, M. Niederweis, H. A. Stenmark, T. H. Flo, Plasma membrane damage causes NLRP3 activation and pyroptosis during Mycobacterium tuberculosis infection. Nat Commun 11, 2270 (2020).

73. C. Bedia, L. Camacho, J. L. Abad, G. Fabriàs, T. Levade, A simple fluorogenic method for determination of acid ceramidase activity and diagnosis of Farber disease. J Lipid Res 51, 3542–3547 (2010).

74. K. R. Brandvold, J. M. Weaver, C. Whidbey, A. T. Wright, A continuous fluorescence assay for simple quantification of bile salt hydrolase activity in the gut microbiome. Sci Rep 9, 1359 (2019).

75. D. I. Benjamin, A. Cozzo, X. Ji, L. S. Roberts, S. M. Louie, M. M. Mulvihill, K. Luo, D. K. Nomura, Ether lipid generating enzyme AGPS alters the balance of structural and signaling lipids to fuel cancer pathogenicity. Proc Natl Acad Sci U S A 110, 14912–14917 (2013).

76. M. Ito, T. Kurita, K. Kita, A novel enzyme that cleaves the N-acyl linkage of ceramides in various glycosphingolipids as well as sphingomyelin to produce their lyso forms. J Biol Chem 270, 24370–24374 (1995).

77. J. P. Koelmel, N. M. Kroeger, E. L. Gill, C. Z. Ulmer, J. A. Bowden, R. E. Patterson, R. A. Yost, T. J. Garrett, Expanding Lipidome Coverage Using LC-MS/MS Data-Dependent Acquisition with Automated Exclusion List Generation. J Am Soc Mass Spectrom 28, 908– 917 (2017).

78. J. D. Ewald, G. Zhou, Y. Lu, J. Kolic, C. Ellis, J. D. Johnson, P. E. Macdonald, J. Xia, Web-based multi-omics integration using the Analyst software suite. Nat Protoc, doi: 10.1038/s41596-023-00950-4 (2024).

79. L. Coulier, R. Bas, S. Jespersen, E. Verheij, M. J. van der Werf, T. Hankemeier, Simultaneous quantitative analysis of metabolites using ion-pair liquid chromatography-electrospray ionization mass spectrometry. Anal Chem 78, 6573–6582 (2006).

80. S. Tyanova, T. Temu, P. Sinitcyn, A. Carlson, M. Y. Hein, T. Geiger, M. Mann, J. Cox, The Perseus computational platform for comprehensive analysis of (prote)omics data. Nat Methods 13, 731–740 (2016).

81. A. J. O’Donoghue, A. A. Eroy-Reveles, G. M. Knudsen, J. Ingram, M. Zhou, J. B. Statnekov, A. L. Greninger, D. R. Hostetter, G. Qu, D. A. Maltby, M. O. Anderson, J. L. Derisi, J. H. McKerrow, A. L. Burlingame, C. S. Craik, Global identification of peptidase specificity by multiplex substrate profiling. Nat Methods 9, 1095–1100 (2012).

82. N. Colaert, K. Helsens, L. Martens, J. Vandekerckhove, K. Gevaert, Improved visualization of protein consensus sequences by iceLogo. Nat Methods 6, 786–787 (2009).

83. F. da Veiga Leprevost, S. E. Haynes, D. M. Avtonomov, H.-Y. Chang, A. K. Shanmugam, D. Mellacheruvu, A. T. Kong, A. I. Nesvizhskii, Philosopher: a versatile toolkit for shotgun proteomics data analysis. Nat Methods 17, 869–870 (2020).

84. A. T. Kong, F. V. Leprevost, D. M. Avtonomov, D. Mellacheruvu, A. I. Nesvizhskii, MSFragger: ultrafast and comprehensive peptide identification in mass spectrometry-based proteomics. Nat Methods 14, 513–520 (2017).

85. A. I. Nesvizhskii, A. Keller, E. Kolker, R. Aebersold, A statistical model for identifying proteins by tandem mass spectrometry. Anal Chem 75, 4646–4658 (2003).

86. G. C. Teo, D. A. Polasky, F. Yu, A. I. Nesvizhskii, Fast Deisotoping Algorithm and Its Implementation in the MSFragger Search Engine. J Proteome Res 20, 498–505 (2021).

87. K. L. Yang, F. Yu, G. C. Teo, K. Li, V. Demichev, M. Ralser, A. I. Nesvizhskii, MSBooster: improving peptide identification rates using deep learning-based features. Nat Commun 14, 4539 (2023).

88. F. Yu, S. E. Haynes, A. I. Nesvizhskii, IonQuant Enables Accurate and Sensitive Label-Free Quantification With FDR-Controlled Match-Between-Runs. Mol Cell Proteomics 20, 100077 (2021).

89. D. J. Abraham, A. J. Leo, Extension of the fragment method to calculate amino acid zwitterion and side chain partition coefficients. Proteins 2, 130–152 (1987).

90. Cancer Genome Atlas Research Network, J. N. Weinstein, E. A. Collisson, G. B. Mills, K. R. M. Shaw, B. A. Ozenberger, K. Ellrott, I. Shmulevich, C. Sander, J. M. Stuart, The Cancer Genome Atlas Pan-Cancer analysis project. Nat Genet 45, 1113–1120 (2013).

91. Gene Ontology Consortium, S. A. Aleksander, J. Balhoff, S. Carbon, J. M. Cherry, H. J. Drabkin, D. Ebert, M. Feuermann, P. Gaudet, N. L. Harris, D. P. Hill, R. Lee, H. Mi, S. Moxon, C. J. Mungall, A. Muruganugan, T. Mushayahama, P. W. Sternberg, P. D. Thomas, K. Van Auken, J. Ramsey, D. A. Siegele, R. L. Chisholm, P. Fey, M. C. Aspromonte, M. V. Nugnes, F. Quaglia, S. Tosatto, M. Giglio, S. Nadendla, G. Antonazzo, H. Attrill, G. Dos Santos, S. Marygold, V. Strelets, C. J. Tabone, J. Thurmond, P. Zhou, S. H. Ahmed, P. Asanitthong, D. Luna Buitrago, M. N. Erdol, M. C. Gage, M. Ali Kadhum, K. Y. C. Li, M. Long, A. Michalak, A. Pesala, A. Pritazahra, S. C. C. Saverimuttu, R. Su, K. E. Thurlow, R. C. Lovering, C. Logie, S. Oliferenko, J. Blake, K. Christie, L. Corbani, M. E. Dolan, H. J. Drabkin, D. P. Hill, L. Ni, D. Sitnikov, C. Smith, A. Cuzick, J. Seager, L. Cooper, J. Elser, P. Jaiswal, P. Gupta, P. Jaiswal, S. Naithani, M. Lera-Ramirez, K. Rutherford, V. Wood, J. L. De Pons, M. R. Dwinell, G. T. Hayman, M. L. Kaldunski, A. E. Kwitek, S. J. F. Laulederkind, M. A. Tutaj, M. Vedi, S.-J. Wang, P. D’Eustachio, L. Aimo, K. Axelsen, A. Bridge, N. Hyka-Nouspikel, A. Morgat, S. A. Aleksander, J. M. Cherry, S. R. Engel, K. Karra, S. R. Miyasato, R. S. Nash, M. S. Skrzypek, S. Weng, E. D. Wong, E. Bakker, T. Z. Berardini, L. Reiser, A. Auchincloss, K. Axelsen, G. Argoud-Puy, M.-C. Blatter, E. Boutet, L. Breuza, A. Bridge, C. Casals-Casas, E. Coudert, A. Estreicher, M. Livia Famiglietti, M. Feuermann, A. Gos, N. Gruaz-Gumowski, C. Hulo, N. Hyka-Nouspikel, F. Jungo, P. Le Mercier, D. Lieberherr, P. Masson, A. Morgat, I. Pedruzzi, L. Pourcel, S. Poux, C. Rivoire, S. Sundaram, A. Bateman, E. Bowler-Barnett, H. Bye-A-Jee, P. Denny, A. Ignatchenko, R. Ishtiaq, A. Lock, Y. Lussi, M. Magrane, M. J. Martin, S. Orchard, P. Raposo, E. Speretta, N. Tyagi, K. Warner, R. Zaru, A. D. Diehl, R. Lee, J. Chan, S. Diamantakis, D. Raciti, M. Zarowiecki, M. Fisher, C. James-Zorn, V. Ponferrada, A. Zorn, S. Ramachandran, L. Ruzicka, M. Westerfield, The Gene Ontology knowledgebase in 2023. Genetics 224, iyad031 (2023).

92. M. Uhlen, C. Zhang, S. Lee, E. Sjöstedt, L. Fagerberg, G. Bidkhori, R. Benfeitas, M. Arif, Z. Liu, F. Edfors, K. Sanli, K. von Feilitzen, P. Oksvold, E. Lundberg, S. Hober, P. Nilsson, J. Mattsson, J. M. Schwenk, H. Brunnström, B. Glimelius, T. Sjöblom, P.-H. Edqvist, D. Djureinovic, P. Micke, C. Lindskog, A. Mardinoglu, F. Ponten, A pathology atlas of the human cancer transcriptome. Science 357, eaan2507 (2017).

93. H. Mi, A. Muruganujan, X. Huang, D. Ebert, C. Mills, X. Guo, P. D. Thomas, Protocol Update for large-scale genome and gene function analysis with the PANTHER classification system (v.14.0). Nat Protoc 14, 703–721 (2019).

94. P. D. Thomas, D. Ebert, A. Muruganujan, T. Mushayahama, L.-P. Albou, H. Mi, PANTHER: Making genome-scale phylogenetics accessible to all. Protein Sci 31, 8–22 (2022).

95. L. Badea, V. Herlea, S. O. Dima, T. Dumitrascu, I. Popescu, Combined gene expression analysis of whole-tissue and microdissected pancreatic ductal adenocarcinoma identifies genes specifically overexpressed in tumor epithelia. Hepatogastroenterology 55, 2016– 2027 (2008).

96. K. A. Ellsworth, B. W. Eckloff, L. Li, I. Moon, B. L. Fridley, G. D. Jenkins, E. Carlson, A. Brisbin, R. Abo, W. Bamlet, G. Petersen, E. D. Wieben, L. Wang, Contribution of FKBP5 genetic variation to gemcitabine treatment and survival in pancreatic adenocarcinoma. PLoS One 8, e70216 (2013).

97. S. Yang, P. He, J. Wang, A. Schetter, W. Tang, N. Funamizu, K. Yanaga, T. Uwagawa, A. R. Satoskar, J. Gaedcke, M. Bernhardt, B. M. Ghadimi, M. M. Gaida, F. Bergmann, J. Werner, T. Ried, N. Hanna, H. R. Alexander, S. P. Hussain, A Novel MIF Signaling Pathway Drives the Malignant Character of Pancreatic Cancer by Targeting NR3C2. Cancer Res 76, 3838–3850 (2016).

98. M. Ghandi, F. W. Huang, J. Jané-Valbuena, G. V. Kryukov, C. C. Lo, E. R. McDonald, J. Barretina, E. T. Gelfand, C. M. Bielski, H. Li, K. Hu, A. Y. Andreev-Drakhlin, J. Kim, J. M. Hess, B. J. Haas, F. Aguet, B. A. Weir, M. V. Rothberg, B. R. Paolella, M. S. Lawrence, R. Akbani, Y. Lu, H. L. Tiv, P. C. Gokhale, A. de Weck, A. A. Mansour, C. Oh, J. Shih, K. Hadi, Y. Rosen, J. Bistline, K. Venkatesan, A. Reddy, D. Sonkin, M. Liu, J. Lehar, J. M. Korn, D. A. Porter, M. D. Jones, J. Golji, G. Caponigro, J. E. Taylor, C. M. Dunning, A. L. Creech, A. C. Warren, J. M. McFarland, M. Zamanighomi, A. Kauffmann, N. Stransky, M. Imielinski, Y. E. Maruvka, A. D. Cherniack, A. Tsherniak, F. Vazquez, J. D. Jaffe, A. A. Lane, D. M. Weinstock, C. M. Johannessen, M. P. Morrissey, F. Stegmeier, R. Schlegel, W. C. Hahn, G. Getz, G. B. Mills, J. S. Boehm, T. R. Golub, L. A. Garraway, W. R. Sellers, Next-generation characterization of the Cancer Cell Line Encyclopedia. Nature 569, 503– 508 (2019).

99. P. Shannon, A. Markiel, O. Ozier, N. S. Baliga, J. T. Wang, D. Ramage, N. Amin, B. Schwikowski, T. Ideker, Cytoscape: a software environment for integrated models of biomolecular interaction networks. Genome Res 13, 2498–2504 (2003).

